# A Bayesian Approach for Detecting Mass-Extinction Events When Rates of Lineage Diversification Vary

**DOI:** 10.1101/020149

**Authors:** Michael R. May, Sebastian Höhna, Brian R. Moore

## Abstract

The paleontological record chronicles numerous episodes of mass extinction that severely culled the Tree of Life. Biologists have long sought to assess the extent to which these events may have impacted particular groups. We present a novel method for detecting mass-extinction events from phylogenies estimated from molecular sequence data. We develop our approach in a Bayesian statistical framework, which enables us to harness prior information on the frequency and magnitude of mass-extinction events. The approach is based on an episodic stochastic-branching process model in which rates of speciation and extinction are constant between rate-shift events. We model three types of events: (1) instantaneous tree-wide shifts in speciation rate; (2) instantaneous tree-wide shifts in extinction rate, and; (3) instantaneous tree-wide mass-extinction events. Each of the events is described by a separate compound Poisson process (CPP) model, where the waiting times between each event are exponentially distributed with event-specific rate parameters. The magnitude of each event is drawn from an event-type specific prior distribution. Parameters of the model are then estimated using a reversible-jump Markov chain Monte Carlo (rjMCMC) algorithm. We demonstrate via simulation that this method has substantial power to detect the number of mass-extinction events, provides unbiased estimates of the timing of mass-extinction events, while exhibiting an appropriate (*i.e.,* below 5%) false discovery rate even in the case of background diversification rate variation. Finally, we provide an empirical application of this approach to conifers, which reveals that this group has experienced two major episodes of mass extinction. This new approach—the CPP on Mass Extinction Times (`CoMET`) model—provides an effective tool for identifying mass-extinction events from molecular phylogenies, even when the history of those groups includes more prosaic temporal variation in diversification rate.

---

The paleontological record documents numerous episodes of mass extinction that severely culled the Tree of Life. As biologists, we often wish to assess whether these events may have impacted particular groups. To this end, several statistical phylogenetic approaches have been proposed to identify mass-extinction events from estimated molecular phylogenies with divergence times, *e.g.,* Nee et al. (1994) and Harvey et al. (1994). These methods generally assume that the dated phylogeny is known without error, and decompose the tree into a vector of waiting times between speciation events. Various models of lineage diversification are then fit to these phylogenetic ‘observations’ using maximum likelihood to estimate the speciation, *b*, and extinction, *d*, rates, and to identify tree-wide shifts in diversification rate, including episodes of mass extinction.

This research program was initially derailed when it was demonstrated that it is not possible to distinguish among different histories of temporal variation in diversification rates within a maximum-likelihood framework (Kubo and Iwasa 1995): an infinite number of unique diversification histories may give rise to an identical vector of waiting times between speciation events in this tree. For this reason, recent developments of inferring diversification process parameters have focused on speciation and extinction rates only (*e.g.,* Rabosky 2006; Paradis 2011; Stadler 2011a; Morlon et al. 2011; Etienne and Haegeman 2012; Höhna 2014). Unfortunately, more realistic diversification models—such as a birth-death process with speciation- and extinction-rate shifts and mass-extinction events—are non-identifiable when parameters are estimated in a maximum-likelihood framework (Stadler 2009; 2011a).

In order to distinguish more prosaic temporal variation in diversification rate from *bona fide* mass-extinction events, we adopt a Bayesian statistical framework that enables us to leverage prior information on the frequency and magnitude of mass-extinction events. Specifically, we develop an episodic stochastic-branching process model where rates of speciation and extinction are constant between rate-shift events. The events are of three types: (1) instantaneous tree-wide shifts in speciation rate; (2) instantaneous tree-wide shifts in extinction rate, and; (3) instantaneous tree-wide mass-extinction events. Each event type is described by a separate compound Poisson process (CPP) model, where the waiting times between events are exponentially distributed with event-specific Poisson-rate parameters. These rate parameters—*λ*_𝔹_, *λ*_𝔹_, and *λ*_𝕄_—therefore control the frequency of speciation-rate shifts, extinction-rate shifts, and mass-extinction events, respectively. When an event occurs, its magnitude is described by the corresponding prior probability density. Specifically, when an event entails a shift in speciation or extinction rate, we draw a new rate from a lognormal prior distribution with hyperpriors *μ* and *σ* specifying the mean and standard deviation, respectively, of the lognormal diversification-rate priors. Similarly, when an event involves an episode of mass extinction, the number of surviving lineages is drawn from a beta prior to describe the survival probability, with hyperpriors *α* and *β* specifying the shape of the beta prior. Between events, the tree evolves under a (piece-wise) constant birth-death stochastic-branching process, motivating its description as an episodic birth-death process.

In principle, all of the free parameters of the CPP on Mass Extinction Time (

~~~
CoMET
~~~

) model could be estimated, including: (1) the number and timing of tree-wide shifts in speciation rate; (2) the number and timing of tree-wide shifts in extinction rate; (3) the rate of speciation and extinction between each rate-shift event, and; (4) the number, timing and severity of tree-wide mass-extinction events. In practice, however, it may not be possible to reliably estimate all of the 

~~~
CoMET
~~~

 model parameters. This limitation stems from the tendency of the CPP model to be non-(or weakly) identifiable (Rannala 2002). For example, when used as a model describing shifts in substitution rate across branches of a phylogeny (the ‘CPP relaxed-molecular clock model’; Huelsenbeck et al. 2000; Blanquart and Lartillot 2008), the CPP model can explain the substitution-rate variation in a given dataset equally well by specifying relatively frequent substitution-rate shifts of small magnitude, or by specifying less frequent substitution-rate shifts of greater magnitude. In fact, there are an infinite number of CPP model parameterizations for which the data have an identical likelihood (*i.e.*, for which the model is non-identifiable). Accordingly, this model is known to be very sensitive to the choice of priors that specify the frequency and magnitude of substitution-rate shifts (*e.g.*, Rannala 2002; Ronquist et al. 2012).

Our use of the CPP model to describe mass-extinction events, however, benefits from the ability to impose strongly informative, empirically grounded priors on the magnitude (survival probability) and frequency (expected number) of mass-extinction events. The former prior derives from the definition of a mass-extinction event as the loss of a specified fraction of species diversity, and the latter can be guided by paleontological information regarding the likely number of mass-extinction events in the relevant period. By contrast, it is difficult to justify strongly informative priors related to the frequency and magnitude of temporal shifts in rates of speciation and extinction. Accordingly, estimates of the corresponding 

~~~
CoMET
~~~

 parameters (model components 1 − 3, above) are expected to be extremely sensitive to the choice of prior, and therefore difficult to estimate reliably. Nevertheless, it is well established that temporal variation in diversification rates is a pervasive feature of empirical phylogenies (*e.g.*, Moen and Morlon 2014), which may impact our ability to accurately identify mass-extinction events. Accordingly, several components (1 − 3) of the 

~~~
CoMET
~~~

 model are included as nuisance parameters intended to improve estimation of the focal parameters: the number and timing mass-extinction events.

In this paper, we first provide a detailed description of the 

~~~
CoMET
~~~

 model and the numerical algorithms used to estimate the parameters of this model. We then perform a comprehensive simulation study to explore the statistical behavior of the 

~~~
CoMET
~~~

 model, including its liability to detect spurious mass-extinction events (the false discovery rate), the ability of the method to detect actual mass-extinction events (the power), and the accuracy of the method to estimate the timing of mass-extinction events (the bias). Finally, we apply our approach to an empirical dataset to reveal the impact of mass-extinction events in coniferous land plants.

## METHODS

### The episodic reconstructed process with explicit mass-extinction events

Our approach is based on the *reconstructed evolutionary process* described by Nee et al. (1994); a birth-death process in which only surviving lineages are observed. Let *N* (*t*) denote the number of species at time *t*. Assume the process starts at time *t*_1_ (the ‘crown’ age of the most recent common ancestor of the study group, *t*_MR__CA_) when there are two species. Thus, the process is initiated with two species, *N* (*t*_1_) = 2. We condition the process on sampling at least one descendant from each of these initial two lineages; otherwise *t*_1_ would not correspond to the *t*_MRCA_ of our study group. Each lineage evolves independently of all other lineages, giving rise to exactly one new lineage with rate *b*(*t*) and losing one existing lineage with rate *d*(*t*) (Figure 1A). Note that although each lineage evolves independently, all lineages share both a common (tree-wide) speciation rate *b*(*t*) and a common extinction rate *d*(*t*) (Nee et al. 1994; Stadler 2011a; Höhna 2013; 2014; 2015). Additionally, at certain times, *t*_𝕄_, a mass-extinction event occurs and each species existing at that time has the same probability, *ρ*, of surviving the mass-extinction event. Finally, all extinct lineages are pruned and only the reconstructed tree remains.

**Figure 1:**
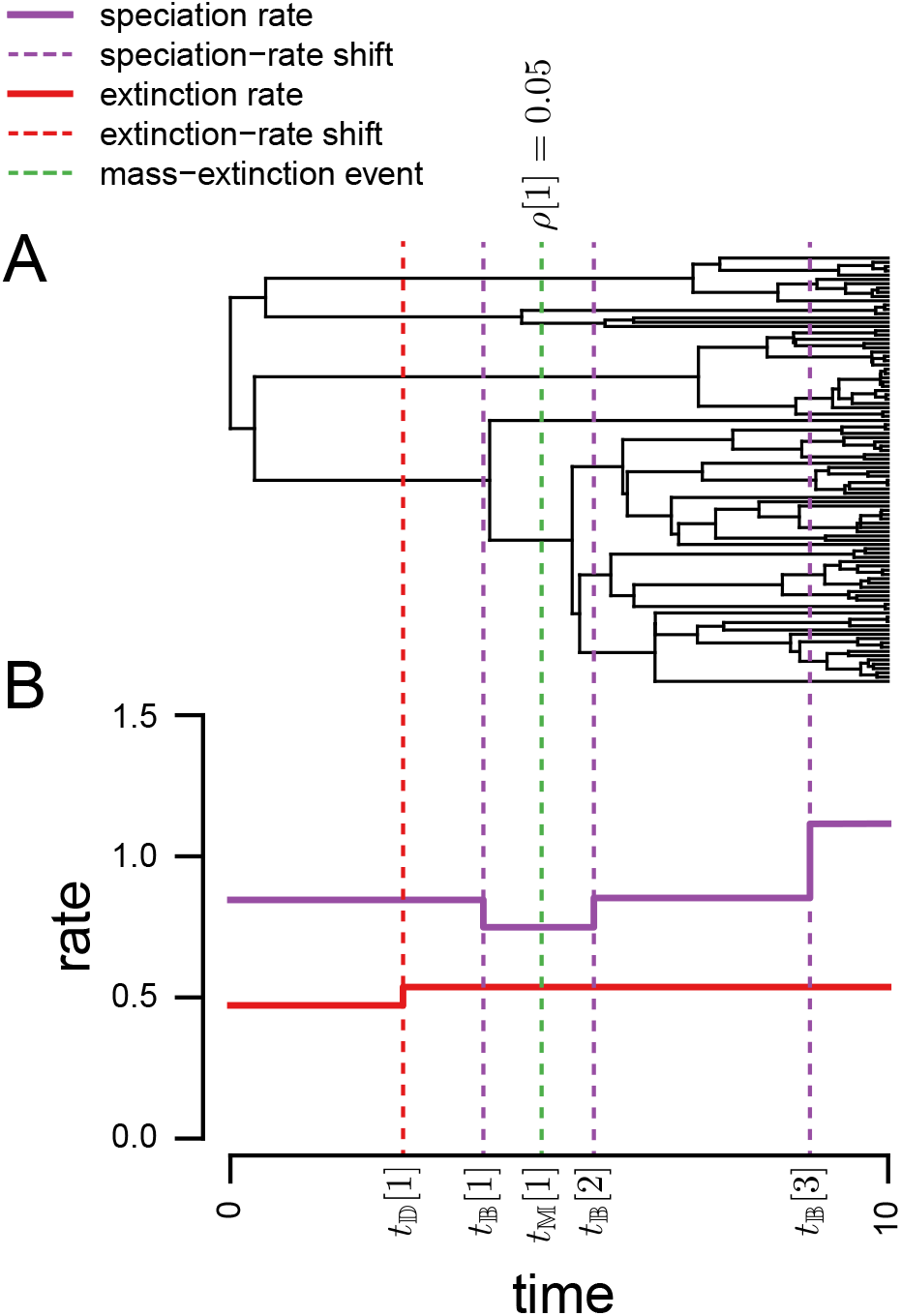
The piecewise-constant birth-death process with mass extinction. A) A realization of the process involves one speciation-rate shift, three extinction-rate shifts, and one mass-extinction event. B) Corresponding plots of the episodic (piecewise constant) speciation and extinction rates, with the times of the five events (see Table 1 for notation). The survival probability for the single mass-extinction event is *ρ* = 0.05.

Our derivation of the 

~~~
CoMET
~~~

 model assumes piecewise-constant speciation and extinction rates that shift instantaneously at rate-change events (see Figure 1B). Therefore, we specify the times of the *k*_𝔹_ speciation-rate shifts in the vector 𝕋_𝔹_ = {*t*_𝔹_[1], …, *t*_𝔹_[*k*_𝔹_]}. We specify the speciation rate within each of the *k* intervals in the vector B = {*b*_0_*, …, b*_*k*_B }, and define the speciation-rate function as *b*(*t*) = *b*_*i*_ for the interval *t*_𝔹_[*i*] *≤ t < t*_𝔹_[*i* + 1]. Similarly, we specify the times of the *k*_𝔹_ extinction-rate shifts in the vector 𝕋_𝔹_ = {*t*_𝔹_[1]*, …, t*_𝔹_[*k*_𝔹_]}. We specify the extinction rates in the vector D = {*d*_0_*, …, d*_*k*_D }, and the extinction-rate function is defined as *d*(*t*) = *d*_*i*_ for the interval *t*_𝔹_[*i*] *≤ t < t*_𝔹_[*i* + 1]. Finally, we specify the times of the *k*_𝕄_ mass-extinction events in the vector 𝕋_𝕄_ = {*t*_𝕄_[1]*, …, t*_𝕄_[*k*_𝕄_]}, where the survival probability for each event is specified in the vector ℙ = {*ρ*_1_*, …, ρ*_*k*_M }. We note that mass-extinction events might be modeled *implicitly*, where a shift to a relatively high extinction rate is followed—after a short interval—by a return to a relatively low extinction rate. By contrast, we model episodes of mass extinction *explicitly* —as instantaneous events—so that we can estimate the probability that such events have occurred.

We present a graphical model description of the 

~~~
CoMET
~~~

 model (Figure S1), and summarize the notation and interpretation of the model parameters in Table 1.

**Table 1:**
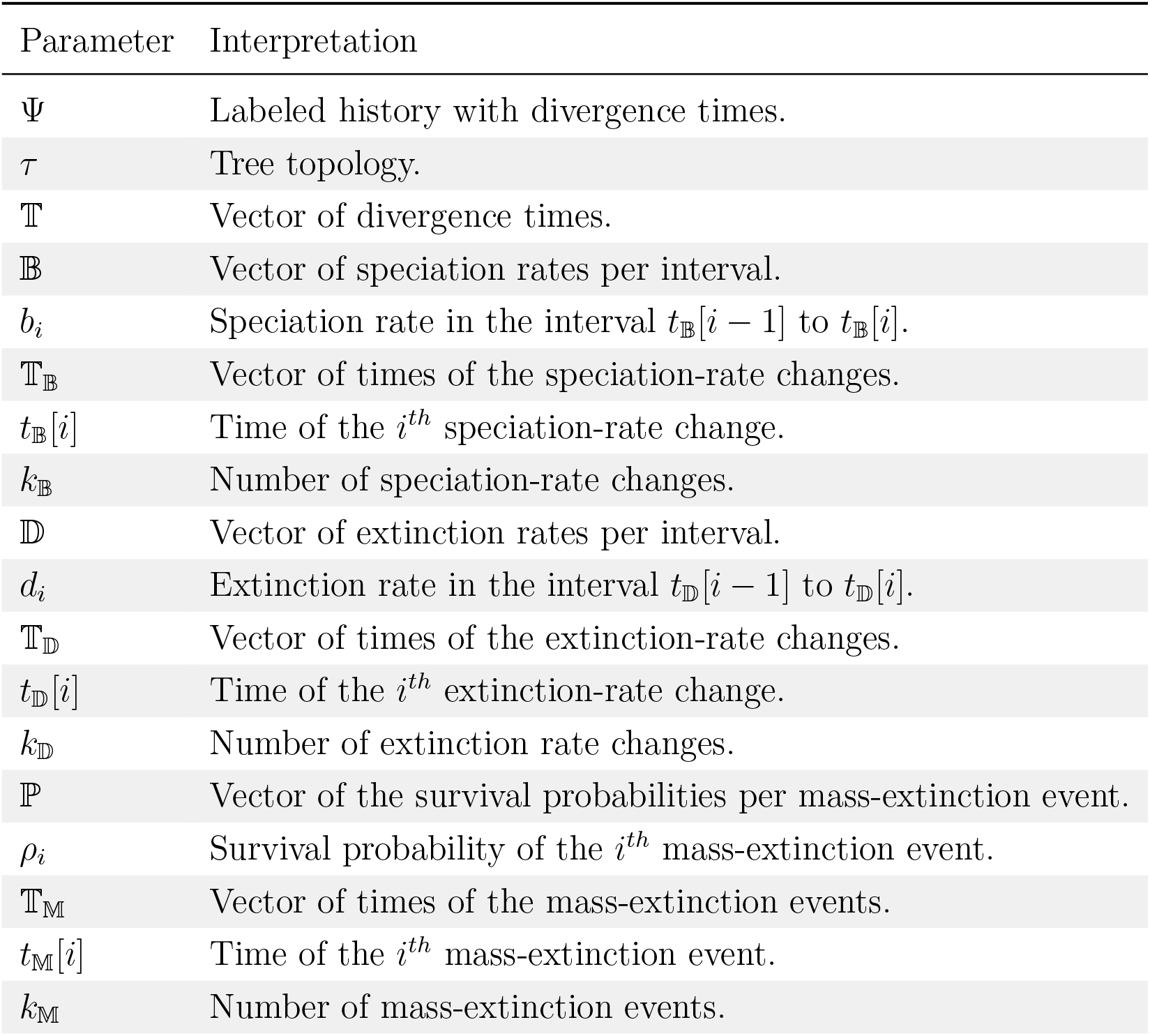
~~~
CoMET
~~~

| Parameter | Interpretation |
| --- | --- |
| $\Psi$ | Labeled history with divergence times. |
| $\tau$ | Tree topology. |
| $\mathbb{T}$ | Vector of divergence times. |
| $\mathbb{B}$ | Vector of speciation rates per interval. |
| $b_i$ | Speciation rate in the interval $t_{\mathbb{B}}[i - 1]$ to $t_{\mathbb{B}}[i]$. |
| $\mathbb{T}_{\mathbb{B}}$ | Vector of times of the speciation-rate changes. |
| $t_{\mathbb{B}}[i]$ | Time of the $i^{th}$ speciation-rate change. |
| $k_{\mathbb{B}}$ | Number of speciation-rate changes. |
| $\mathbb{D}$ | Vector of extinction rates per interval. |
| $d_i$ | Extinction rate in the interval $t_{\mathbb{D}}[i - 1]$ to $t_{\mathbb{D}}[i]$. |
| $\mathbb{T}_{\mathbb{D}}$ | Vector of times of the extinction-rate changes. |
| $t_{\mathbb{D}}[i]$ | Time of the $i^{th}$ extinction-rate change. |
| $k_{\mathbb{D}}$ | Number of extinction rate changes. |
| $\mathbb{P}$ | Vector of the survival probabilities per mass-extinction event. |
| $\rho_i$ | Survival probability of the $i^{th}$ mass-extinction event. |
| $\mathbb{T}_{\mathbb{M}}$ | Vector of times of the mass-extinction events. |
| $t_{\mathbb{M}}[i]$ | Time of the $i^{th}$ mass-extinction event. |
| $k_{\mathbb{M}}$ | Number of mass-extinction events. |

 **model parameters and their interpretation**

*Likelihood function.*—Let ψ denote a reconstructed evolutionary tree relating *n* species, comprising a tree topology, *τ*, and the set of branching times, 𝕋. For a birth-death process where the rates of speciation and extinction are the same for all branches at any instant in time, the probabilities of the tree topology and the branching times are independent. Thus, we can compute the probability of the reconstructed evolutionary tree as the product of the independent probabilities: *P* (ψ) = *P* (*τ*) *× P* (𝕋).

For a tree with *n* species, there are *n*!(*n* − 1)!*/*2^*n-*1^ unique labeled histories (we use labeled histories and tree topologies interchangeably). Under a birth-death process, all labeled histories are equally likely (Edwards 1970; Rannala and Yang 1996), so

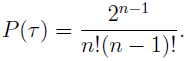

We call a lineage that begins with a single species at time *t* and ends with a single sampled species at time *T* (the present) a *singleton lineage*. We proceed by recognizing that a reconstructed evolutionary tree is composed of a set of independently evolving singleton lineages: a tree with a single node (the root) has two singleton lineages, and each additional node generates an additional singleton lineage (Figure 2A). Under a birth-death process, the probability density of a singleton lineage is just the probability of starting with a single species at time *t* and ending with a single sampled lineage at the present *T*, *P* (*N* (*T*) = 1 *| N* (*t*) = 1) (Figure 2B).

**Figure 2.**
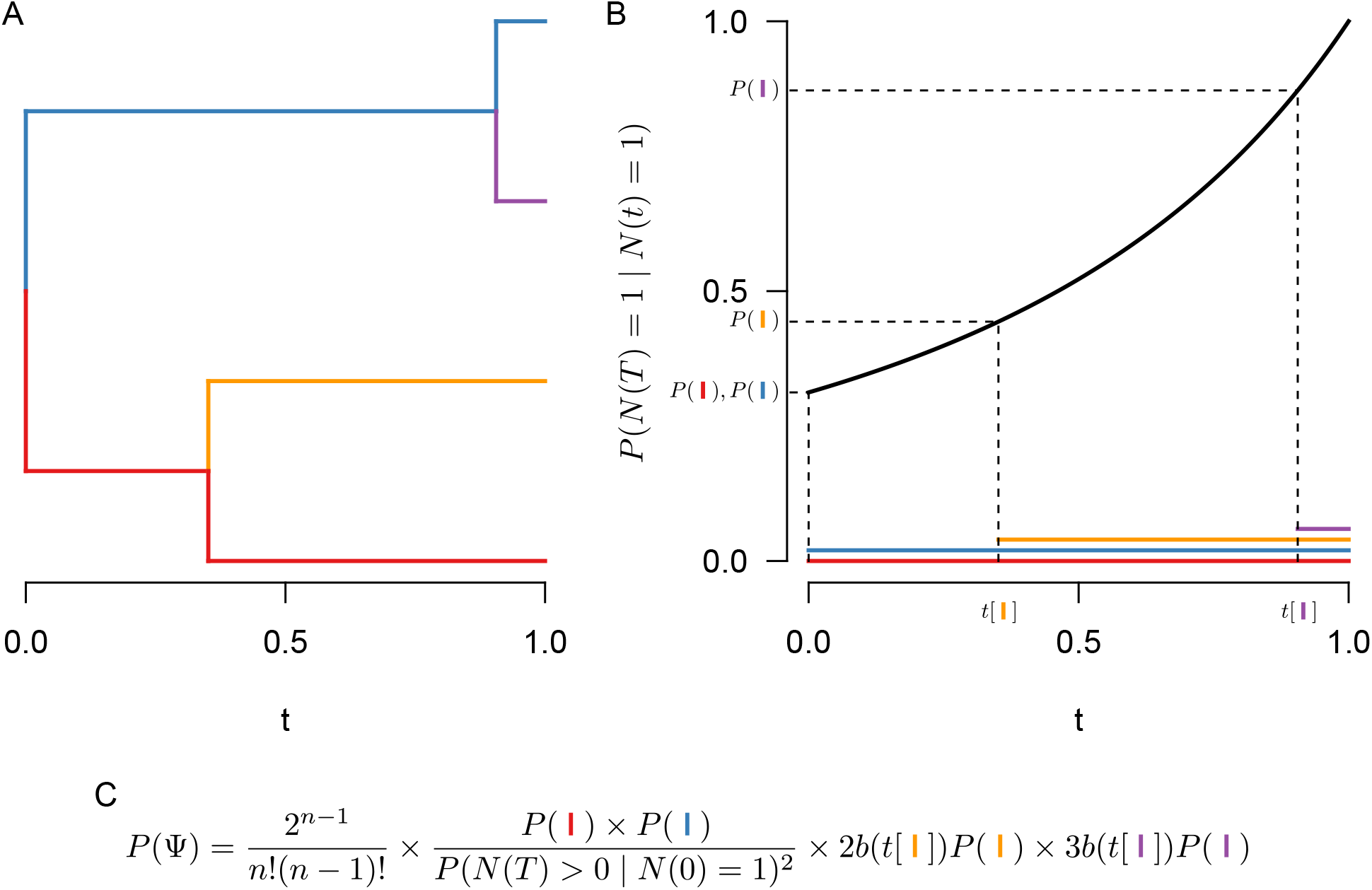
Computing the likelihood of a phylogeny under the birth-death process. We first identify lineages within the phylogeny that begin and end with a single species (panel A, colored branches). Then, for each of those lineages, we compute the probability that the lineage ended with a single extant species at time *T* given that it began with a single species at time *t*, *P* (*N* (*T*) = 1 *N* (*t*) = 1) (panel B). We then multiply those probabilities by the number of labeled histories, as well as the probability that there were speciation events at each non-root node in the tree, *b*(*t*); finally, we condition on survival of the process by dividing by the probability that each lineage descending from the root left at least one extant descendant, *P* (*N* (*T*) *>* 0 *| N* (0) = 1)^2^ (panel C, see also Equation (1)).

We must also incorporate the probability density that each new singleton lineage arises in the first place (*i.e.,* that there is a speciation event at time *t*). Each singleton lineage gives rise to new species at rate *b*(*t*); therefore, in general, the probability density that a speciation event occurs at time *t* is simply *b*(*t*) multiplied by the number of singleton lineages that exist at *t*. For the episodic model we have described, *b*(*t*) = *b*_*i*_ for the interval *t*_𝔹_[*i*] *≤ t < t*_𝔹_[*i* + 1].

We condition the probability density of observing the branching times on the survival of both lineages that descend from the root (otherwise, the root would not exist). To do so, we divide by *P* (*N* (*T*) *>* 0*|N* (0) = 1)^2^.

The probability density of the branching times, 𝕋, becomes

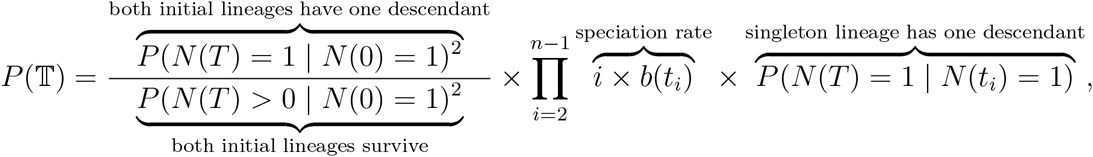

and the probability density of the reconstructed tree (topology and branching times) is then

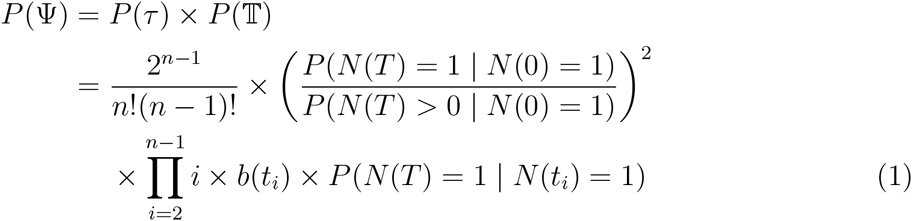

We can simplify Equation (1) by substituting *P* (*N* (*T*) *>* 0 *N* (*t*) = 1)^2^ exp(*r*(*t, T*)) for *P* (*N* (*T*) = 1 *| N* (*t*) = 1), where 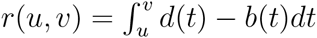 (for a detailed description of this substitution, see Höhna 2015); the above equation becomes

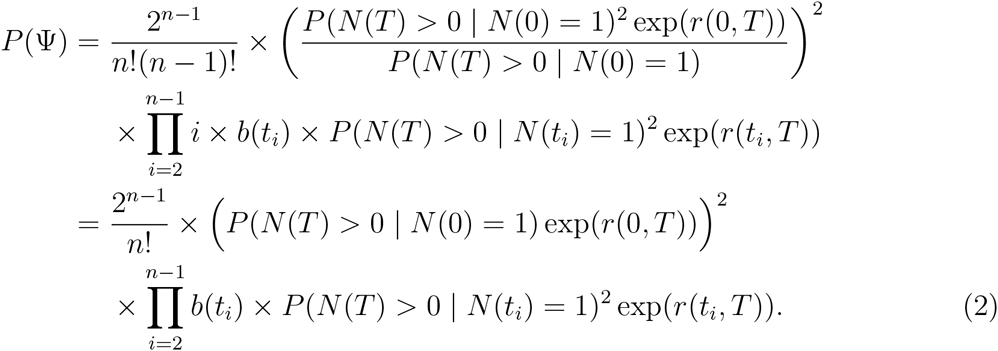

This probability density was originally derived by Thompson (Thompson 1975; Equation (3.4.6)) for constant rates (see also Equation 20 in Nee et al. 1994) and later extended to arbitrary rate functions (Lambert 2010; Höhna 2013; 2014; 2015).

The probability density of a reconstructed phylogeny ψ in Equation (2) is given for any time-dependent birth-death process. Analytical solutions to this equation can be obtained if the following quantities can be computed analytically: *b*(*t*_*i*_), *P* (*N* (*T*) *>* 0*|N* (*t*_*i*_) = 1), and *r*(*t*_*i*_*, T*). Given the speciation and extinction rate and the mass-extinction survival probabilities, we can compute the probability of no extinction (see Equation 16 in Höhna 2015)

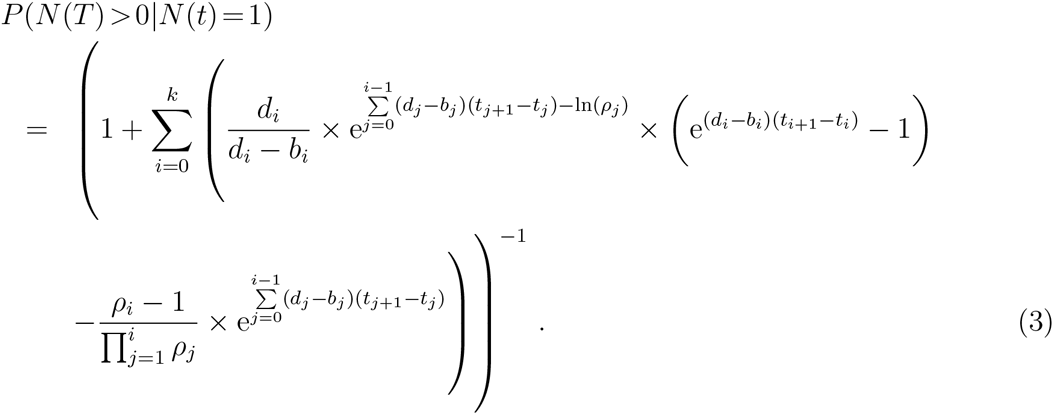

Inserting Equation (3) into Equation (2) yields the probability density of an observed (*i.e.,* reconstructed) tree under the episodic birth-death process with explicit mass-extinction events. We provide this expanded equation in the Appendix; see Equation (4).

### Bayesian Inference

*Parameterization and prior distributions.*—In the previous section we described the episodic birth-death process with mass-extinction events and gave the probability density of an observed tree given the parameters, *i.e.,* the likelihood function for the 

~~~
CoMET
~~~

 model. The likelihood function allows us to estimate parameters of the model using different statistical approaches, including maximum-likelihood estimation and Bayesian inference. Previously, the study of temporal variation in diversification rates has largely been pursued in a maximum-likelihood framework (*e.g.,* Rabosky 2006; Stadler 2011a; Höhna 2014). However, Kubo and Iwasa (1995) demonstrated that stochastic-branching process models are non-identifiable when parameters are estimated using maximum likelihood. That is, the phylogenetic observations (the vector of waiting times between speciation events) are equally likely to be the outcome of an infinite number of distinct diversification processes. For example, a diversification process in which a low initial speciation rate later shifts to a higher speciation rate produces the same phylogenetic observations as a constant-rate process with a mass-extinction event (Stadler 2011b).

These considerations motivated our adoption of a Bayesian solution to this problem. Pursuing the detection of mass-extinction events within a Bayesian statistical framework both allows us to specify a prior distribution on the number of events—thereby automatically penalizing more complex histories—and also to leverage biologically relevant information (as informative priors) on the survival probability of mass-extinction events. Specifically, we draw the number of speciation and extinction rate-shifts from a Poisson prior with rate *λ*_𝔹_ and *λ*_𝔹_, respectively. Following a shift in speciation or extinction rate, we draw a new rate from a lognormal prior with parameters *μ*_𝔹_ and *σ*_𝔹_ or *μ*_𝔹_ and *σ*_𝔹_. Similarly, we draw the number of mass-extinction events from a Poisson prior with rate *λ*_𝕄_, and draw the survival probability from a Beta prior with parameters *α* and *β*. By default, we assume that *α* = 2 and *β* = 18; this corresponds to a prior belief that a mass-extinction event will on average result in the loss 90% of the contemporaneous species diversity. Accordingly, the parameters and prior densities of the 

~~~
CoMET
~~~

 model are as follows:

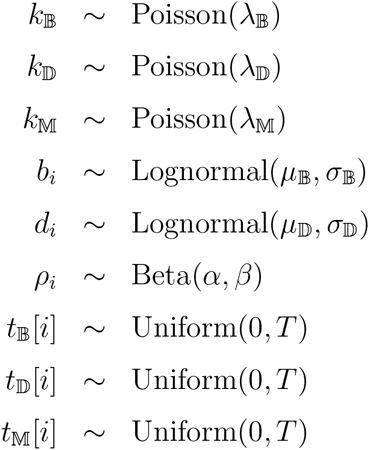

*Empirical Bayesian hyperpriors.*—Our use of lognormal priors for the speciation and extinction rates raises the issue of how we should parameterize these distributions. Specifically, we need to specify the mean and standard deviation of these prior densities. Were we to specify a lognormal prior that is too narrow (*i.e.,* where the standard deviation is too small), the rates would be close to the mean and the model would disfavor large rate shifts. Conversely, if we were to specify a lognormal prior that is too diffuse (*i.e.,* where the standard deviation is too large), the rates would be overly dispersed and the model would tend to overfit patterns of rate variation the data.

It is therefore crucial to carefully select the parameters of these prior distributions. We might pursue one of three possible approaches for specifying the prior mean and standard deviation of the lognormal priors: (1) we could adopt an ‘empirical prior’ approach that treats the mean and standard deviation as fixed values, perhaps guided by biological information on these parameters; (2) we could adopt a ‘hierarchical Bayesian’ approach that treats the mean and standard deviation as random variables, allowing these parameters to be estimated from the data (Holder and Lewis 2003), or; (3) we could adopt an ‘empirical Bayesian’ approach, where the values of these parameters are guided by the data at hand (*c.f.*, Huelsenbeck and Bollback 2001; Yang et al. 2005).

Use of empirical priors is not viable because we typically lack information regarding reasonable values for the mean and standard deviation of speciation and extinction rates. The development of a hierarchical Bayesian model would provide an elegant solution, but defers rather than solves the problem. That is, we avoid specifying values for the mean and standard deviation of the lognormal priors by treating them as random variables, but this necessitates that we specify both the type of second-order hyperpriors (beta, gamma, etc.) and the associated second-order hyperparameters (shape, scale, etc.) for these variables. There are also some immediate practical concerns with a hierarchical Bayesian solution. First, this solution involves adding an extra level of complexity to the 

~~~
CoMET
~~~

 model, which may complicate our characterization of the statistical behavior of this new method. Additionally, a hierarchical Bayesian solution is more computationally demanding, which—although not an issue for empirical applications of the 

~~~
CoMET
~~~

 model—is of more concern for our simulation study that involves a large number of analyses.

Accordingly, we have adopted an empirical Bayesian approach for parameterizing the lognormal priors for the speciation- and extinction-rate parameters. This involves performing a preliminary MCMC simulation for the data at hand under a constrained 

~~~
CoMET
~~~

 model—where rates of speciation and extinction are assumed to be constant—to estimate the posterior probability densities for the speciation and extinction rates. We then center the lognormal prior on the inferred posterior mean for each parameter. Similarly, we specify the standard deviation of the lognormal prior such that the variance of the prior density is ten-fold that of the inferred posterior density.

*Markov chain Monte Carlo implementation.*—We approximate the posterior probability distribution of the 

~~~
CoMET
~~~

 model parameters using a Metropolis-Hastings algorithm (Metropolis et al. 1953; Hastings 1970; Gelman et al. 2003). Specifically, we employ a Markov chain Monte Carlo (MCMC) simulation where we propose updates for all numeric parameters using normally distributed proposal densities centered on the current values (Gelman et al. 2003; Yang and Rodrίguez 2013), and propose updates for the number of events using reversible-jump MCMC (rjMCMC; Green 1995; 2003). We implemented two rjMCMC proposals to add or remove an event—the ‘birth move’ and ‘death move’, respectively—following Huelsenbeck et al. (2000).

#### Birth move

1. Simulate the time of the new event: *t*_*k*+1_ ∼ unif(0*, T*)
2. Simulate the parameter value for the new event from the corresponding prior: *θ*_*k*+1_ ∼ Prior, such that *θ*^*′*^ = {*θ ∪ θ*_*k*+1_}
3. Compute the posterior probability for the proposed value: *f* (*θ*^*′*^) *∞* Likelihood(*θ*^*′*^) *×* Prior(*θ*^*′*^)
4. Compute the posterior probability for the current value: *f* (*θ*) *∝* Likelihood(*θ*) *×* Prior(*θ*)
5. Compute the forward proposal probability: *q*(*θ*^*′*^*|θ*) = 1*/T ×* Prior(*θ*_*k*+1_)
6. Compute the reverse proposal probability: *q*(*θ|θ*^*′*^) = 1*/*(*k* + 1)
7. Compute the Jacobian: J = 1
8. Compute the acceptance probability: *α* = *f* (*θ*^*′*^)*/f* (*θ*) *× q*(*θ|θ*^*′*^)*/q*(*θ*^*′*^*|θ*) *× J* = *f* (*θ*^*′*^)*/f* (*θ*) *×* (1/*T ×* Prior(*θ*_*k*+1_))^−1^

#### Death move

1. Select an event to delete: *idx* ∼ unif(1*, k*) so that *θ*^*′*^ = {*θ\θ*_*idx*_}
2. Compute the posterior probability for the proposed value: *f* (*θ*^*′*^) *∞* Likelihood(*θ*^*′*^) *×* Prior(*θ*^*′*^)
3. Compute the posterior probability for the current value: *f* (*θ*) *∝* Likelihood(*θ*) *×* Prior(*θ*)
4. Compute the forward proposal probability: *q*(*θ*^*′*^*|θ*) = 1/*k*
5. Compute the reverse proposal probability: *q*(*θ|θ*^*′*^) = 1*/T ×* Prior(*θ*_*idx*_) *×* 1/*k*
6. Compute the Jacobian: J = 1
7. Compute the acceptance probability: *α* = *f* (*θ*^*′*^)*/f* (*θ*) *× q*(*θ|θ*^*′*^)*/q*(*θ*^*′*^*|θ*) *× J* = *f* (*θ*^*′*^)*/f* (*θ*) *×* 1/*T ×* Prior(*θ*_*idx*_)

Both birth and death proposals are used to update the number of speciation rate-shifts, extinction rate-shifts, and mass-extinction events. When the number of events is being updated, a birth move or a death move will be applied with equal probability during an iteration of the MCMC simulation. We validated the algorithms and our implementation by sampling from the prior distribution. When the rjMCMC simulation is run without data, it will target the joint *prior* probability density of the model parameters. This allows us to compare the *inferred* marginal prior probability density to the corresponding *known* prior probability density for each model parameter: if the rjMCMC algorithm and implementation are correct, we will recover the known prior densities. These experiments confirmed the validity of the 

~~~
CoMET
~~~

 algorithms.

The 

~~~
CoMET
~~~

 model and the rjMCMC algorithm are implemented in the 

~~~
R
~~~

 package 

~~~
TESS
~~~

 and are available from http://cran.r-project.org/.

### Hypothesis Testing

*Testing hypotheses regarding the timing of significant mass-extinction events.*— Explicitly modeling mass-extinction events enables us to perform robust Bayesian hypothesis testing using Bayes factors. The Bayes factor compares the relative performance of two models (denoted *M*_0_ and *M*_1_) by comparing their *marginal likelihoods*:

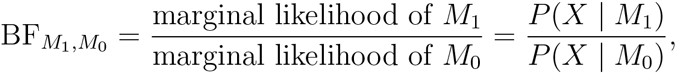

where the marginal likelihood, *P* (*X | M*_*i*_) = *θ P* (*X | θ*)*P* (*θ | M*_*i*_)*dθ*, is the likelihood of the data, *X*, integrated over the entire joint prior distribution of the model parameters (*i.e.,* the average probability of observing the data under the model). Values of BF_*M*_1*,M*0 greater than 1 indicate a preference for *M*_1_, whereas values of BF_*M*_1*,M*0 less than one indicates a preference for *M*_0_ (Kass and Raftery 1995).

Normally, the marginal likelihood is an intractable quantity to compute; however, the Bayes factor can be re-written as:

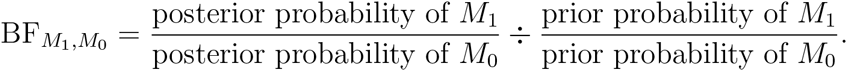

For example, we may be interested in testing the hypothesis that mass extinction *i* occurred at time *t*:
Unfortunately, because we must approximate the posterior probability density of mass-extinction times with MCMC, the (estimated) posterior probability *P* (*t*_𝕄_[*i*] = *t | X*) will always be 0 (*i.e.,* the probability that a numerical sample takes some real value *t* is 0). We can, however, test the hypothesis that at least one mass extinction occurred in the interval *I* = (*t, t* + Δ*t*):

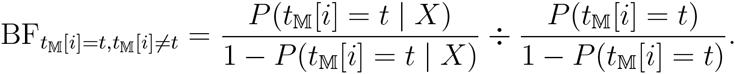

Conveniently, we can calculate the prior probability of no mass extinction under the CPP model:

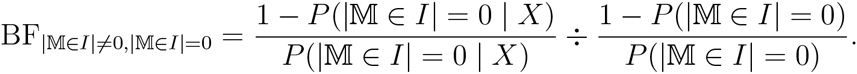

where 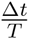 is the duration of the interval relative to the height of the tree. The posterior probability can be approximated directly from the MCMC samples:

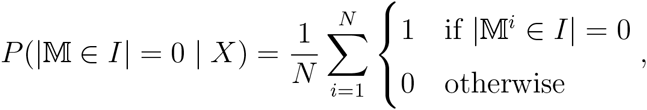

where *N* is the number of MCMC samples and M*i* is the vector of mass-extinction times in the *i*^th^ sample.

Our procedure for identifying the timing of mass-extinction events is as follows: (1) discretize the interval (0*, T*) into *n* intervals of duration 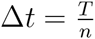; (2) for each interval, compute the Bayes factor for at least one mass extinction in the interval; (3) identify intervals with significant Bayes factor support for the specified significance threshold, BF_crit_, as containing a mass-extinction event, and; (4) merge contiguous runs of intervals with mass-extinction events into a single mass extinction whose time corresponds to the interval with the highest support. This procedure is depicted in Figure 3.

**Figure 3:**
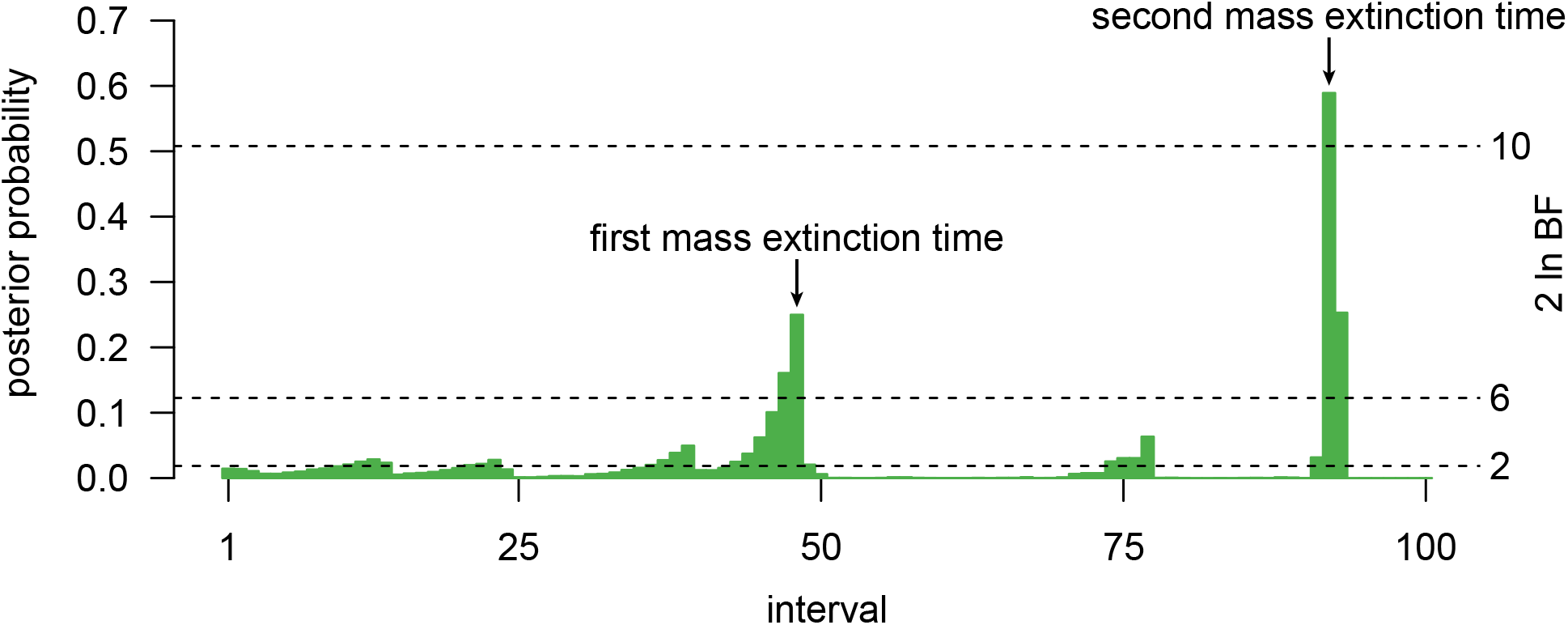
Identifying significant mass-extinction events using Bayes factors. An example of the procedure for estimating the timing of significant mass-extinction events. Each bar represents the posterior probability of at least one mass extinction in that interval. Bars that exceed the significance threshold (in this case, 2 ln BF *>* 6) indicate significant mass extinction events. When multiple adjacent bars are greater than the significance threshold, only the bar with the greatest support is considered a mass extinction. In this example, the Bayes factors are computed assuming *λ*_𝕄_ = ln 2, and we infer significant mass-extinction events in intervals 48 and 93.

There are several practical considerations for this approach. Intervals that are too large relative to *T* will provide imprecise estimates of the mass-extinction times, and may in fact include multiple well-supported mass extinctions, confounding the interpretation of the Bayes factor test. Conversely, intervals that are very small will lead to more precise estimates of mass-extinction times, but will also decrease the number of sampled mass extinctions in the interval, resulting in unstable estimates of the posterior probability. We have found that 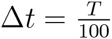 provides a good compromise between precision and stability. The identification of well-supported mass extinctions relies on a significance threshold, BF_crit_. By convention, we use a significance threshold that corresponds to “strong” support (2 ln BF_crit_ *≥* 6, Kass and Raftery 1995). A well-supported mass extinction may also appear in multiple consecutive intervals, which motivates step 4, the merger of contiguous mass extinctions. We note that setting Δ*t* = *T* is equivalent to testing the hypothesis regarding the occurrence of *any* significant mass-extinction events over the entire tree; however, we can also test hypotheses about the *exact number* of mass-extinction events, which we describe in the next section.

*Testing hypotheses regarding the number of significant mass-extinction events.*—Bayes factors can also be used to test hypotheses related to the number of mass-extinction events. By treating the number of mass extinctions as the model, we can assess the support for exactly *k* mass-extinction events:

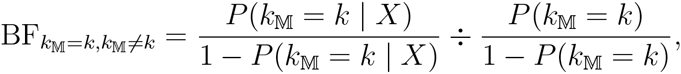

where, under the CPP model, the prior probability is simply calculated as:

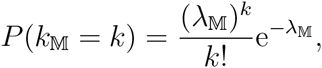

and the posterior probability is directly estimated from the MCMC samples:

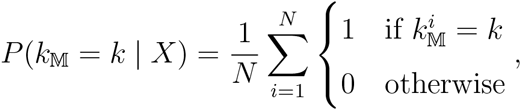

where 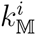 is the number of mass-extinction events in the *i*^th^ MCMC sample.

### Simulation Study

The complex nature of both the 

~~~
CoMET
~~~

 model and the algorithms used to estimate parameters of the model demand a comprehensive simulation study to characterize the statistical behavior of this new method. We designed our simulation study to understand: (1) the rate at which mass-extinction events are incorrectly inferred (the *false discovery rate*); (2) the rate at which mass-extinction events are correctly inferred (the *power*); (3) the accuracy of the inferred timing of mass-extinction events (the *bias*); (4) the ability to distinguish multiple mass-extinction events, and; (5) the influence of shifts in background diversification rates on the false discovery rate, power and bias of our approach. All simulations and analyses were performed in the 

~~~
R
~~~

 package 

~~~
TESS
~~~

 (Höhna 2013).

#### False discovery rate

We first assessed the liability of the 

~~~
CoMET
~~~

 model to detect spurious mass-extinction events in trees simulated under constant speciation and extinction rates. For each tree, we sampled the speciation rate, *b*, from a lognormal distribution with mean *μ*_𝔹_ = 1 and standard deviation *σ*_𝔹_ = exp(0.2). Similarly, we sampled the extinction rate, *d*, from a lognormal distribution with mean *μ*_𝔹_ = 0.5 and standard deviation *σ*_𝔹_ = exp(0.2). We ran each simulation for *T* = 10 time units, generating trees with *N* = {100, 200, 400, 800} species. For each tree size, we simulated 100 trees (400 trees in total; *c.f.*, Figure 4A).

**Figure 4:**
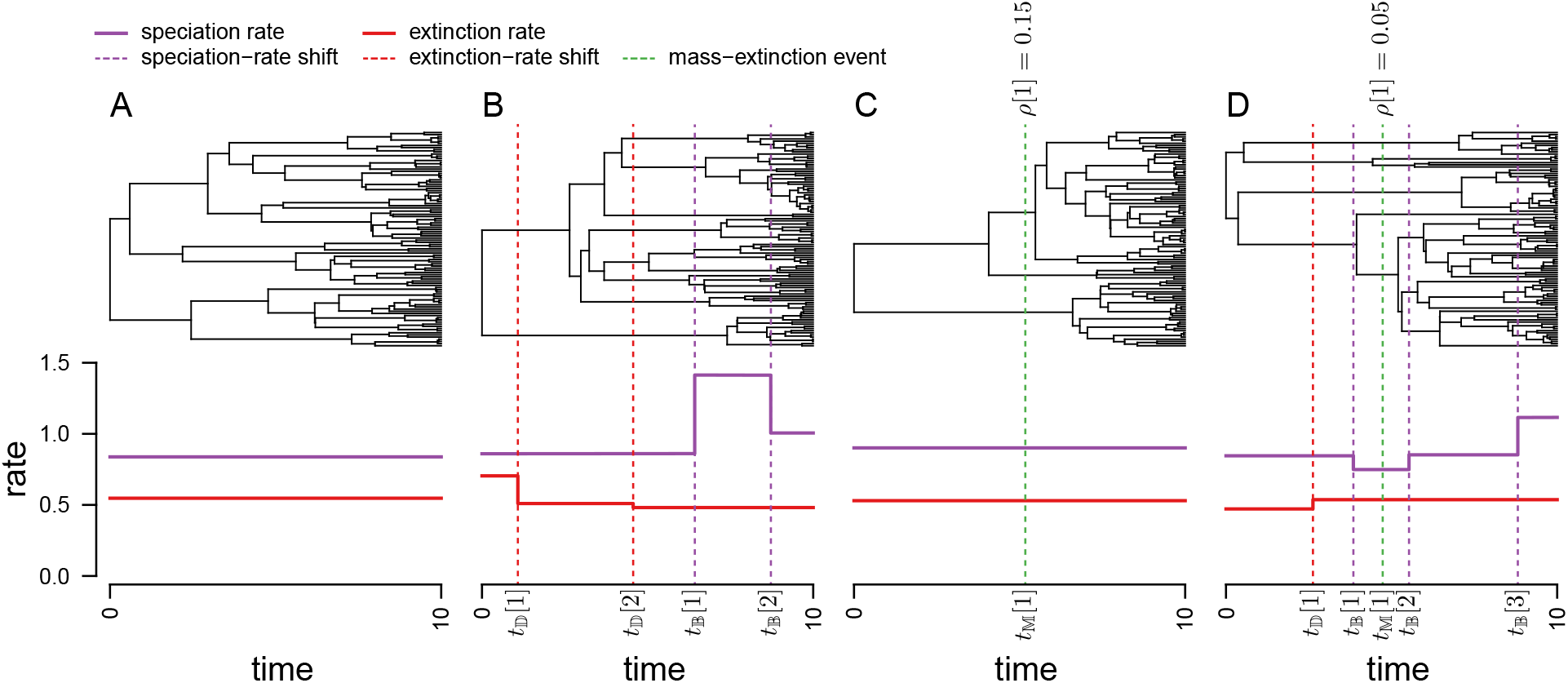
Simulation study design. Panels A and B depict simulations that explore the false discovery rate (where trees are simulated *without* mass-extinction events); panels C and D depict simulations that explore the power and bias (where trees are simulated *with* mass-extinction events). Panels A and C entail simulations where the speciation and extinction rates are constant; panels B and D entail simulations where the speciation and extinction rates shifted episoodically. Each panel shows an actual tree (above) from the simulations with *N* = 100 species, as well as the parameters of the stochastic-branching-process model used in those simulations (below).

We also assessed the liability of the 

~~~
CoMET
~~~

 model to detect spurious mass-extinction events in trees simulated under episodically shifting speciation and extinction rates. For each tree, we sampled the number of speciation- and extinction-rate shifts, *k*_𝔹_ and *k*_𝔹_, from a Poisson distribution with rate parameters *λ*_𝔹_ = *λ*_𝔹_ = 2. We sampled the times of the speciation- and extinction-rate shifts, 𝕋_𝔹_ = {*t*_𝔹_[1], …, *t*_𝔹_[*k*_𝔹_]} and 𝕋_𝔹_ = {*t*_𝔹_[1]*, …, t*_𝔹_[*k*_𝔹_]}, from a uniform distribution on (0*, T*). We sampled the speciation rates, B = {*b*_0_*, …, b*_*k*_B }, from a lognormal distribution with mean *μ*_𝔹_ = 1 and standard deviation *σ*_𝔹_ = exp(0.2). Similarly, we sampled the extinction rates, D = {*d*_0_*, …, d*_*k*_D }, from a lognormal distribution with mean *μ*_𝔹_ = 0.5 and standard deviation *σ*_𝔹_ = exp(0.2). We ran each simulation for *T* = 10 time units, simulating 100 trees of each size, with *N* = {100, 200, 400, 800} species (400 trees in total; *c.f.*, Figure 4B).

In order to explore the impact of the chosen priors on our ability to detect mass-extinction events, we analyzed each simulated tree under a variety of prior settings. We considered cases where the prior expected relative-extinction rate (*d ÷ b*) was either too low, centered on the correct value, or too high. We achieved this by varying the hyperprior on the mean extinction rate, *μ*_𝔹_ = {0.1, 0.5, 0.9}, while fixing the other hyperpriors to the generating values. Specifically, we set the mean and standard deviation of the lognormal speciation-rate prior to *μ*_𝔹_ = 1 and *σ*_𝔹_ = exp(0.2), respectively, and set the standard deviation of the lognormal extinction-rate prior to *σ*_𝔹_ = exp(0.2). In addition to analyses using fixed hyperprior values, we also performed analyses where the values of the hyperpriors for the speciation and extinction rates were estimated using the empirical Bayesian approach described above. We also varied the priors on the frequency of diversification-rate shifts, *λ*_𝔹_ = *λ*_𝔹_ = {0.1, ln 2, 2, 5}, and the frequency of mass-extinction events, *λ*_𝕄_ = {0.1, ln 2, 2, 5}. We set the hyperpriors on the mass-extinction survival probability to *α* = 2, *β* = 18. We analyzed each simulated tree under every combination of prior settings, resulting in 4 *×* 4 *×* 4 = 64 analyses per tree, for a total of 64 *×* 800 = 51, 200 MCMC analyses.

We ran each MCMC simulation until one of two stopping conditions was reached: (1) the effective sample size (ESS, computed with the 

~~~
R
~~~

 package coda) for all of the event-rate parameters—*k*_𝔹_, *k*_𝔹_ and *k*_𝕄_—exceeded 500, or; (2) the maximum number of cycles (one million) was reached. We thinned the chains by sampling every 100^th^ state. Occasionally, one or more parameters were found to have low ESS values (*≤* 200) after the MCMC simulation reached the maximum length. In such cases, we repeated the analysis. We discarded the first 25% of the samples for each MCMC simulation as burnin. We then classified any analyses that identified strong support for at least one mass-extinction event as a false positive, and computed the false discovery rate for a particular combination of prior settings as the fraction of analyses that contained false positives.

#### Power

We first assessed the power of the 

~~~
CoMET
~~~

 model to correctly detect mass-extinction events against a background of constant speciation and extinction rates. We sampled speciation and extinction rates as described above for the false discovery rate experiments. For each tree, we sampled the number of mass-extinction events, *k*_𝕄_, from a Poisson distribution with a rate parameter *λ*_𝕄_ = 1. We sampled the times of the mass-extinction events, 𝕋_𝕄_ = {*t*_𝕄_[1], …, *t*_𝕄_[*k*_𝕄_]}, from a uniform distribution on the interval (0*, T*). For each mass-extinction event, we sampled the survival probabilities, ℙ, from a beta distribution with shape parameters *α* = 2 and *β* = 18, so that the expected survival probability 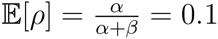. We ran each simulation for *T* = 10 time units, simulating 100 trees of each size, with *N* = {100, 200, 400, 800} species (400 trees in total; *c.f.*, Figure 4C).

We also assessed the power of the 

~~~
CoMET
~~~

 model to correctly identify mass-extinction events against a background of episodically shifting speciation and extinction rates. For this experiment, we simulated trees exactly as in the initial power experiment, except that we simulated diversification-rate shifts as described in the false discovery rate experiment. We ran each simulation for *T* = 10 time units, simulating 100 trees of each size, with *N* = {100, 200, 400, 800} species (400 trees in total; *c.f.*, Figure 4D).

We analyzed each simulated tree under the same variety of prior settings, and performed and diagnosed the MCMC simulations as described in the false discovery rate experiments (51, 800 analyses in total). We classified an analysis as having correctly inferred a mass-extinction event if it indicated strong support for one mass-extinction event when the tree actually experienced one mass-extinction event. We computed the power for a particular combination of prior settings as the fraction of analyses that correctly identified a mass-extinction event.

#### Bias

We assessed the ability of the 

~~~
CoMET
~~~

 model to accurately estimate the timing of mass-extinction events using the analyses from the previous section. When an analysis correctly inferred a single mass-extinction event, we computed the bias of the estimated event time as (*t*_simulated_ − *t*_estimated_)*/*tree height *×* 100%.

#### Multiple mass-extinction events

Motivated by our empirical conifer analysis (see below), we investigated the behavior of the 

~~~
CoMET
~~~

 model under sequential mass-extinction events. In particular, we were interested in the ability of the method to successfully detect the older of two mass-extinction events, and the ability to distinguish the two mass extinctions as a function of their relative age.

To approximate the conifer dataset (see below), we simulated trees with *N* = 492 species and a tree height of *T* = 340.43 million years. We simulated the trees under constant background speciation and extinction rates, which were sampled from the posterior distributions estimated from our empirical analyses. We simulated one ancient mass-extinction event at *t*_a_ = 250 Ma to mimic the Permo-Triassic mass-extinction event. We then simulated a second, more recent mass extinction at *t*_r_ = {200, 150, 100, 50} Ma. Both mass-extinction events had a survival probability of *ρ* = 0.1. We simulated 100 trees for each value of *t*_r_, for a total of 400 simulated trees.

We analyzed each tree using empirical Bayesian estimates of the diversification-rate hyperpriors—*μ*_𝔹_*, σ*_𝔹_ and *μ*_𝔹_*, σ*_𝔹_—and assumed a constant background diversification rate (*λ*_𝔹_ = *λ*_𝔹_ = 0). We analyzed each simulated tree under a variety of prior settings for the frequency of mass-extinction events, *λ*_𝕄_ = {0.1, ln 2, 2, 5}, for a total of 4 *×* 400 = 1, 600 analyses. We performed MCMC analyses and diagnostics as described previously.

For each analysis, we then estimated the time of inferred mass-extinction events as described previously in the hypothesis-testing section. We considered 

~~~
CoMET
~~~

 to have provided a correct result if the *inferred* mass-extinction event was within four intervals [4% = (340.43 *÷* 100) *×* 4 = ±13.6 million years] of the *simulated* mass-extinction event. We then calculated the fraction of analyses under each set of priors that identified the more recent, the more ancient, or both mass-extinction events.

### An Empirical Example: Mass Extinction in Conifers

To demonstrate the application of the 

~~~
CoMET
~~~

 model to an empirical dataset, we present an analysis exploring mass-extinction events in conifers. Our analysis is based on a recent study of the phylogeny and divergence times of conifers (Leslie et al. 2012), which included 492 of 630 (78%) described species and inferred a crown age of 340.43 Ma. Accordingly, this conifer tree spans three major mass-extinction events; the Permo-Triassic (252 Ma), the Triassic-Jurassic (201.3 Ma), and the Cretaceous-Paleogene (66 Ma) mass-extinction events. Each event is estimated to have caused the loss of *∼* 70 − 75% of contemporaneous terrestrial species (*e.g.,* Raup and Sepkoski 1982; Labandeira and Sepkoski 1993; Rees 2002; McElwain and Punyasena 2007; Cascales-Miñana and Cleal 2014).

We conditioned our conifer analyses on the maximum-clade credibility consensus tree from the Leslie et al. (2012) study, with the three cycad (outgroup) species removed. We first performed a series of preliminary analyses on this tree to estimate the marginal posterior probability densities for the diversification-rate hyperpriors, *μ*_𝔹_*, σ*_𝔹_*, μ*_𝔹_, and *σ*_𝔹_. We performed these analyses under a constrained 

~~~
CoMET
~~~

 model, where background diversification rates were held constant and mass-extinction events were precluded (specified by setting *λ*_𝔹_ = *λ*_𝔹_ = *λ*_𝕄_ = 0). We approximated the joint posterior probability density under this constrained model by running four independent MCMC simulations, and thinned each chain by sampling every 1,000^th^ state. The chains were terminated when the ESS values for every parameter reached 500. We discarded the first 25% of samples from each chain as burnin, and combined the stationary samples from the four independent chains. We then used the inferred composite marginal posterior probability densities to specify values for the diversification-rate hyperpriors; that is, for the mean and standard deviation of the lognormal priors on the speciation and extinction rates, *μ*_𝔹_*, σ*_𝔹_*, μ*_𝔹_, and *σ*_𝔹_. Specifically, we centered each of the lognormal priors on the corresponding estimate of the posterior mean, and specified the standard deviation such that the variance of the prior density was ten-fold that of the corresponding inferred marginal posterior density. The inferred marginal posterior densities for the diversification rate hyperparameters—and the marginal hyperprior densities elicited from them—are depicted in Figure S8.

We then performed a second series of analyses under the full 

~~~
CoMET
~~~

 model to infer the history of mass extinction in conifers. For these analyses, we used the previously specified empirical Bayesian diversification-rate hyperpriors (*i.e., μ*_𝔹_*, σ*_𝔹_*, μ*_𝔹_, and *σ*_𝔹_), and assumed phylogenetically uniform species sampling (*e.g.,* Höhna et al. 2011; Höhna 2014). We specified the prior probability of surviving mass-extinction events using a beta prior with shape parameters *α* = 2.5 and *β* = 7.5; this specifies an expected survival probability of 25%, which is consistent with prior knowledge (McElwain and Punyasena 2007; Cascales-Miñana and Cleal 2014). As in our analyses of the simulated datasets, we assessed the prior sensitivity of our inferences by performing analyses under a variety of priors. Specifically, we performed analyses under a range of values for the prior on the frequency of mass-extinction events, *λ*_𝕄_ = {0.1, ln 2, 2, 5}, and also for the priors on the frequency of shifts in diversification rate, *λ*_𝔹_ = *λ*_𝔹_ = {0.1, ln 2, 2, 5}. For each unique combination of prior settings, we approximated the joint posterior probability density by running four independent MCMC simulations, and thinned each chain by sampling every 1,000^th^ state. We terminated terminated an MCMC simulation when the ESS values for every parameter reached 500, discarded the first 25% of samples from each chain as burnin, and then combined the stationary samples from the four independent chains. Our estimates of the number and timing of mass-extinction events were based on the resulting composite marginal posterior probability density for each of the unique prior settings.

#### Data availability statement

The authors confirm that all data supporting the results of this study are fully available without restriction. We have made these data available as an archive—including all of the empirical and simulated datasets, as well as the scripts used to simulate and analyze those data—that has been deposited in the Dryad digital repository. The Dryad data identifier for this archive is: doi:xx.xxxx/dryad.xxxxx.

## RESULTS

### Simulation Study

#### False discovery rates

We evaluated the false discovery rate (FDR) of the 

~~~
CoMET
~~~

 model by computing the fraction of cases where it identified strong support for one or more spurious mass-extinction events in trees that were simulated without episodes of mass extinction. Figure 5 depicts the FDR for analyses of trees with *N* = 400 species, where the extinction rate was centered on the true value (*μ*_𝔹_ = 0.5, left column), or where the diversification-rate hyperpriors were specified using the empirical Bayesian approach (right column). Results were qualitatively similar across all analyses (see Figures S2 and S3).

**Figure 5:**
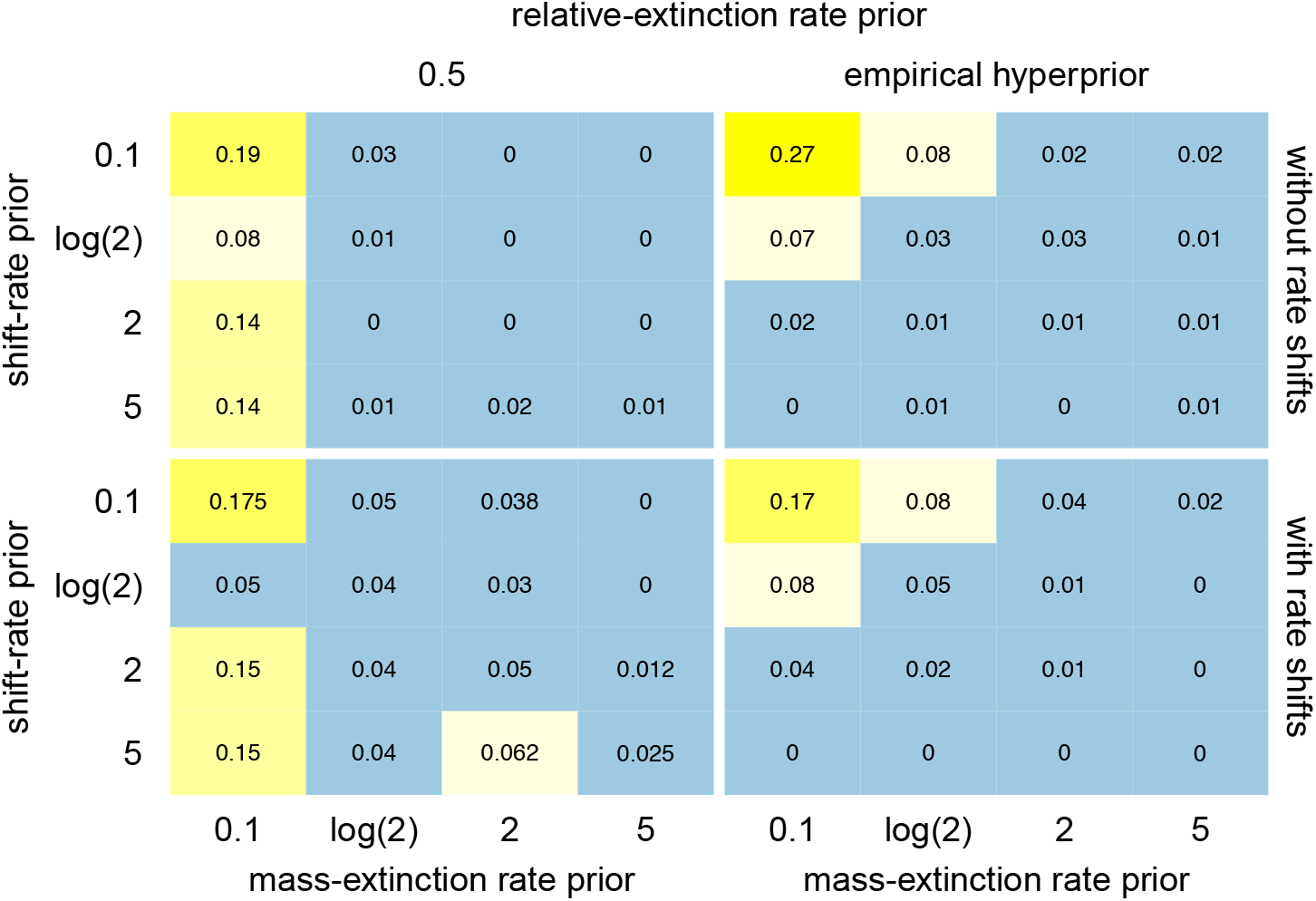
False discovery rates. Frequency of detecting spurious mass-extinction events. Rows of panels correspond to the absence (top panels) or presence (bottom panels) of background diversification-rate shifts, and columns of panels correspond to relative-extinction rate priors centered on the true values (left panels) or estimated from the data (right panels). Within each panel, the rows correspond to false discovery rates under various priors on the expected number of diversification-rate shifts (rows) and mass-extinction events (columns).

Our simulation study implies that the 

~~~
CoMET
~~~

 model has a slightly elevated FDR (relative to the conventional 5% threshold): the overall FDR was inferred to be 8.1% or 9.9% for trees simulated under constant or episodically shifting background diversification rates, respectively. However, a more careful examination of the results reveals that spurious mass-extinction events are far more likely under specific combinations of prior settings. Specifically, the false discovery rate was substantially inflated when the prior rate of mass-extinction events was very low (FDR for *λ*_𝕄_ = 0.1: 18.9% without diversification-rate shifts, 21.2% with diversification-rate shifts), compared to the remainder of the analyses (FDR for *λ*_𝕄_≠ 0.1: 4.4% without diversification-rate shifts, 6.1% with diversification-rate shifts). The prior mean on the extinction rate, *μ*_𝔹_, had a less pronounced effect on the false discovery rate (FDR for *μ*_𝔹_ = {0.1, 0.5, 0.9}: 10.8%, 5.3%, 13.5% without diversification-rate shifts; 12.6%, 7.4%, 15.3% with diversification-rate shifts). Importantly, the false discovery rate was much lower for analyses using empirical Bayesian approach to specify hyperpriors (FDR for empirical hyperprior analyses: 2.8% without diversification-rate shifts, 4.1% with diversification-rate shifts).

#### Power

We evaluated the power of the 

~~~
CoMET
~~~

 model by computing the fraction of cases where it identified strong support for a known (*i.e.*, simulated) mass-extinction event. Moreover, we assessed power as a function of the relative timing of mass-extinction events; we placed each analysis into one of six bins corresponding to the simulated mass-extinction time and computed the power for each of the bins under each combination of prior settings. Figure 6 depicts the statistical power of the 

~~~
CoMET
~~~

 model for trees with *N* = 400 species, where the extinction rate was centered on the true value (*μ*_𝔹_ = 0.5, left column), or where the diversification-rate hyperpriors were specified using the empirical Bayesian approach (right column). Results were qualitatively similar across all analyses (see Figures S4 and S5).

**Figure 6:**
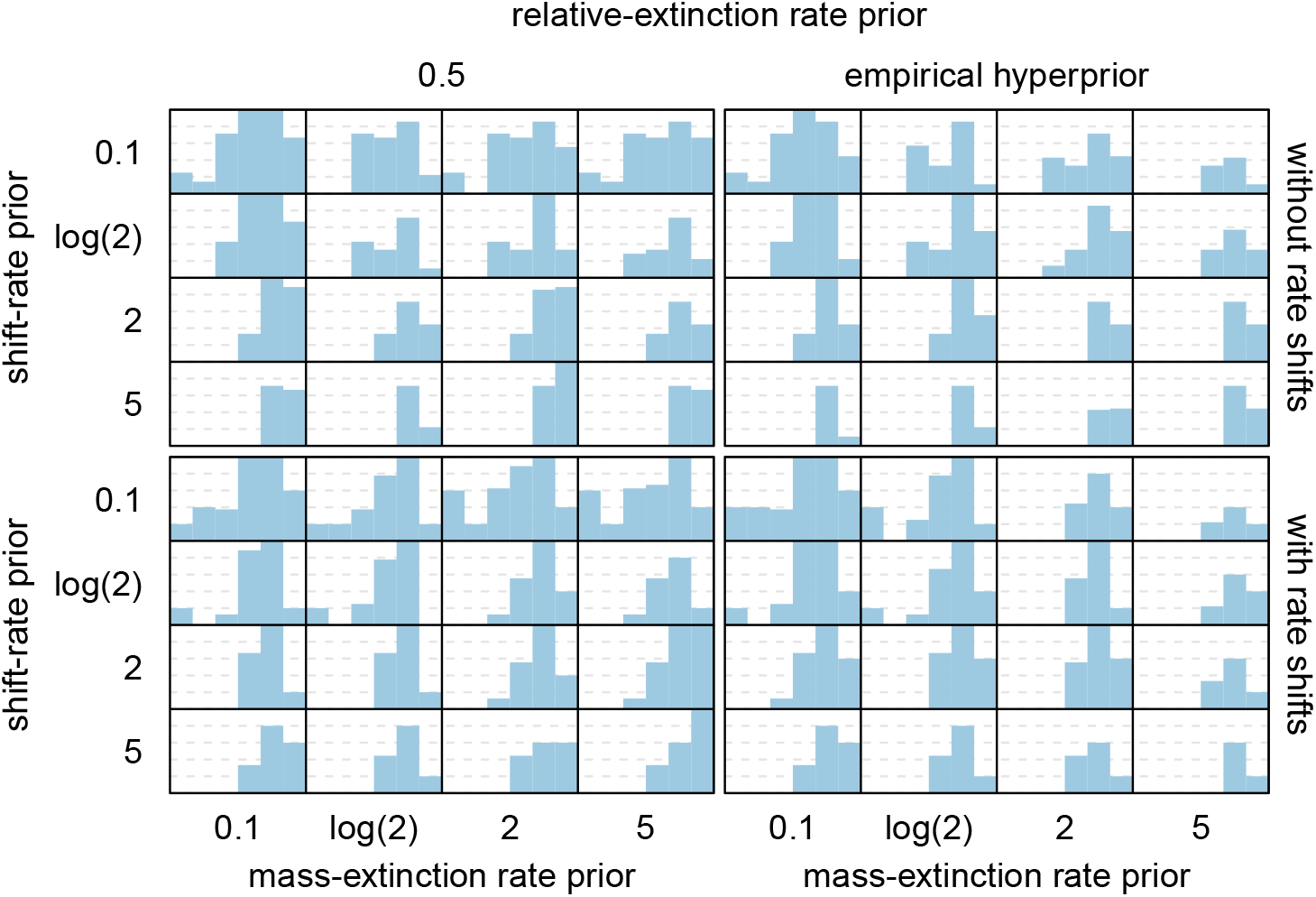
Power as a function of the relative time of mass-extinction events. Frequency of detecting true mass-extinction events. Rows of panels correspond to the absence (top panels) or presence (bottom panels) of background diversification-rate shifts, and columns of panels correspond to relative-extinction rate priors centered on the true values (left panels) or estimated from the data (right panels). Within each panel, the rows correspond to the power under various priors on the expected number of diversification-rate shifts (rows) and mass-extinction events (columns). Within each cell, we plot power as a function of time by binning simulated trees by the relative time of the detected mass-extinction event, and compute the fraction of those trees where a mass-extinction event was correctly inferred.

Our ability to correctly infer mass-extinction events depended critically on the timing of the event. Detection rates were much higher in the more recent half of the tree (power in the recent half: 51.5% without diversification-rate shifts, 53.7% with diversification rate shifts) compared to the more ancient half (power in the ancient half: 8.8% without diversification-rate shifts, 11.6% with diversification rate shifts). Indeed, power was the greatest when the mass-extinction event occurred somewhere between 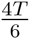 and 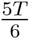 (power: 64.0% without diversification-rate shifts, 58.8% with diversification-rate shifts). The prior on mass-extinction rate had a small influence on the power, with smaller values having greater power (power in the more recent half of the tree for *λ*_𝕄_ = {2, ln 2, 2, 5}: 60.1%, 65.3%, 47.7%, 32.4% without diversification-rate shifts, 59.7%, 65.7%, 49.9%, 39.6% with diversification-rate shifts). Importantly, the empirical Bayes approach did not markedly reduce overall power (power in the more recent half of the tree for empirical hyperpriors: 50.4% without diversification rate-shifts, 54.3% with diversification-rate shifts; compared to *a priori* hyperpriors: 51.7% without diversification-rate shifts, 53.5% with diversification rate-shifts). As expected, overall power increased with the size tree, especially in the face of diversification-rate shifts (power in the more recent half of the tree for *N* = {100, 200, 400, 800}: 47.1%, 54.5%, 48.8%, 55.6% without diversification-rate shifts, 40.6%, 56.2%, 55.9%, 62.1% with diversification rate-shifts).

#### Bias

We evaluated the bias of inferred mass-extinction times under the 

~~~
CoMET
~~~

 model as follows: for all cases in which a mass-extinction event was correctly inferred, we compared the actual event time to the estimated event time as (*t*_actual_ − *t*_estimated_)*/*tree height *×* 100%. Figure 7 depicts the bias of the 

~~~
CoMET
~~~

 model for analyses of trees with *N* = 400 species, where the extinction rate was centered on the true value (*μ*_𝔹_ = 0.5, left column), or where the diversification-rate hyperpriors were specified using the empirical Bayesian approach (right column). Results were similar for the entire simulation (see Figures S6 and S7). Across all analyses, the estimated mass-extinction times were approximately unbiased; overall, the inferred event times were biased by −0.5% or −3.2% of the true event times for trees simulated under constant or episodically shifting background diversification rates, respectively.

**Figure 7:**
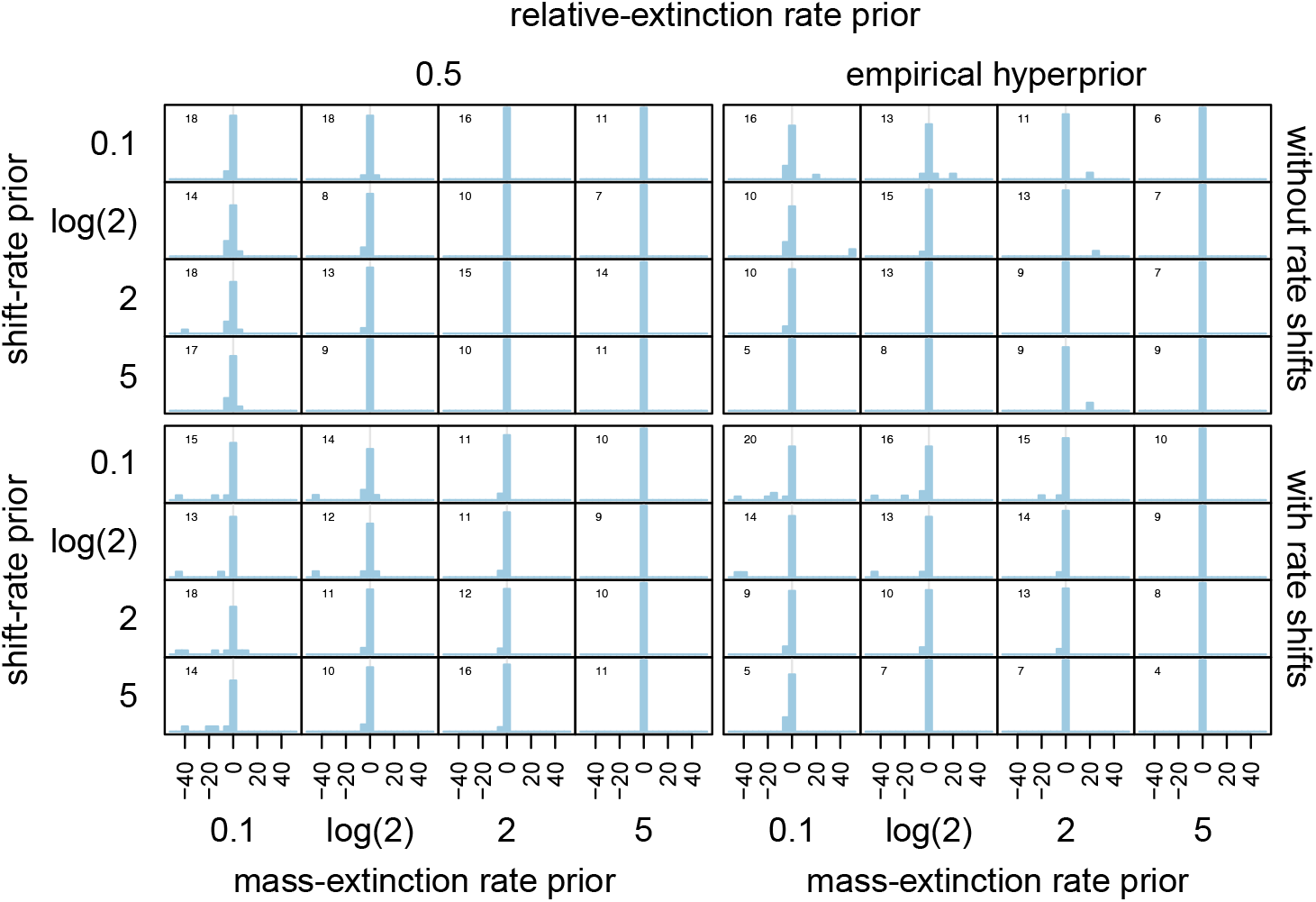
Bias in the estimated timing of mass-extinction events. Rows of panels correspond to the absence (top panels) or presence (bottom panels) of background diversification-rate shifts, and columns of panels correspond to relative-extinction rate priors centered on the true values (left panels) or estimated from the data (right panels). Within each panel, rows correspond to the bias under various priors on the expected number of diversification-rate shifts (rows) and mass-extinction events (columns). We computed the bias as (*t*simulated event − *t*estimated event)*/*tree height *×* 100%.

#### Multiple mass-extinction events

Figure 8 summarizes results of our empirically motivated investigation of sequential mass-extinction events. The ability of the 

~~~
CoMET
~~~

 model to detect the more recent mass extinction depends on the relative age of the event: we detected *≈* 90% of the events that occurred 50 Ma, but only *≈* 15% of those that occurred 200 Ma. These findings are consistent with our results based on the single-event simulations, which indicate that power increases with the recency of the mass-extinction event (*c.f.*, Figure 6). Similarly, our ability to detect the more ancient event depends on the interval between the sequential mass-extinction events: the more recent event casts a shadow backward in time that can completely eclipse an older event if they are spaced too close together in time.

**Figure 8:**
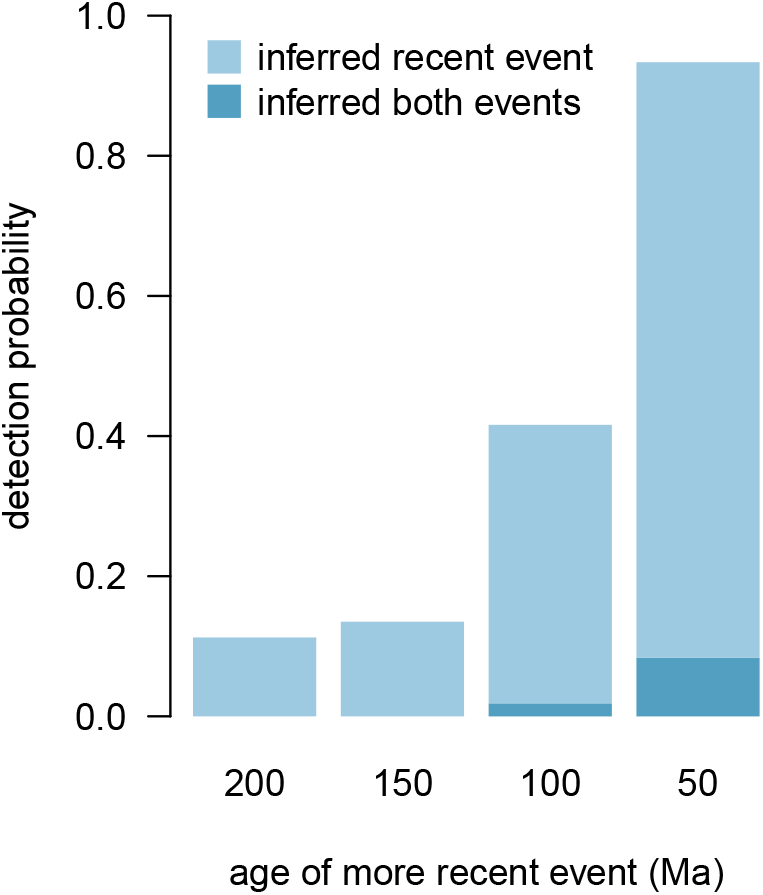
Power to detect sequential mass-extinction events. We plot the frequency of correctly detecting relatively recent (open circles) and ancient (open triangles) mass-extinction events as a function of the age of the more recent event. We plot the age of the more recent mass-extinction event along the x-axis, and the probability of correctly detecting mass-extinction events along the y-axis.

### Mass Extinction in Conifers

Our analysis of the conifer phylogeny identified two mass-extinction events: the first event was inferred to have occurred approximately 173 Ma with positive support, and the second event to have occurred approximately 23 Ma with very strong support (Figure 9). Based on the results of our simulation study, we suspect that the “shadow effect” of the more recent event has diminished the signal of the earlier mass-extinction event. Curiously, the inferred times of the two events do not coincide with the established ages of major mass-extinction events. The closest known events to the inferred mass-extinction times are the Eocene-Oligocene event (34 Ma) and the Toarcian-turnover event (183 Ma); however, these mass-extinction events are not thought to have strongly impacted terrestrial flora (McElwain and Punyasena 2007; Cascales-Miñana and Cleal 2014)

Our estimates of the mass-extinction times in conifers are, of course, based on divergence-time estimates for this group. Accordingly, any bias in the inferred divergence times will cause a corresponding bias in the inferred timing of the mass-extinction events. If, for example, conifer divergence times are slightly underestimated, then the timing of the more recent event could conceivably correspond to the Cretaceous-Paleogene mass-extinction event (66 Ma), and the age of the earlier event could easily correspond to the Triassic-Jurassic (201.3 Ma) or the Permo-Triassic mass-extinction events (252 Ma). We note, however, that our method is unlikely to detect both the Triassic-Jurassic and the Permo-Triassic mass-extinction events, given their close temporal proximity (*c.f.*, Figure 8).

**Figure 9:**
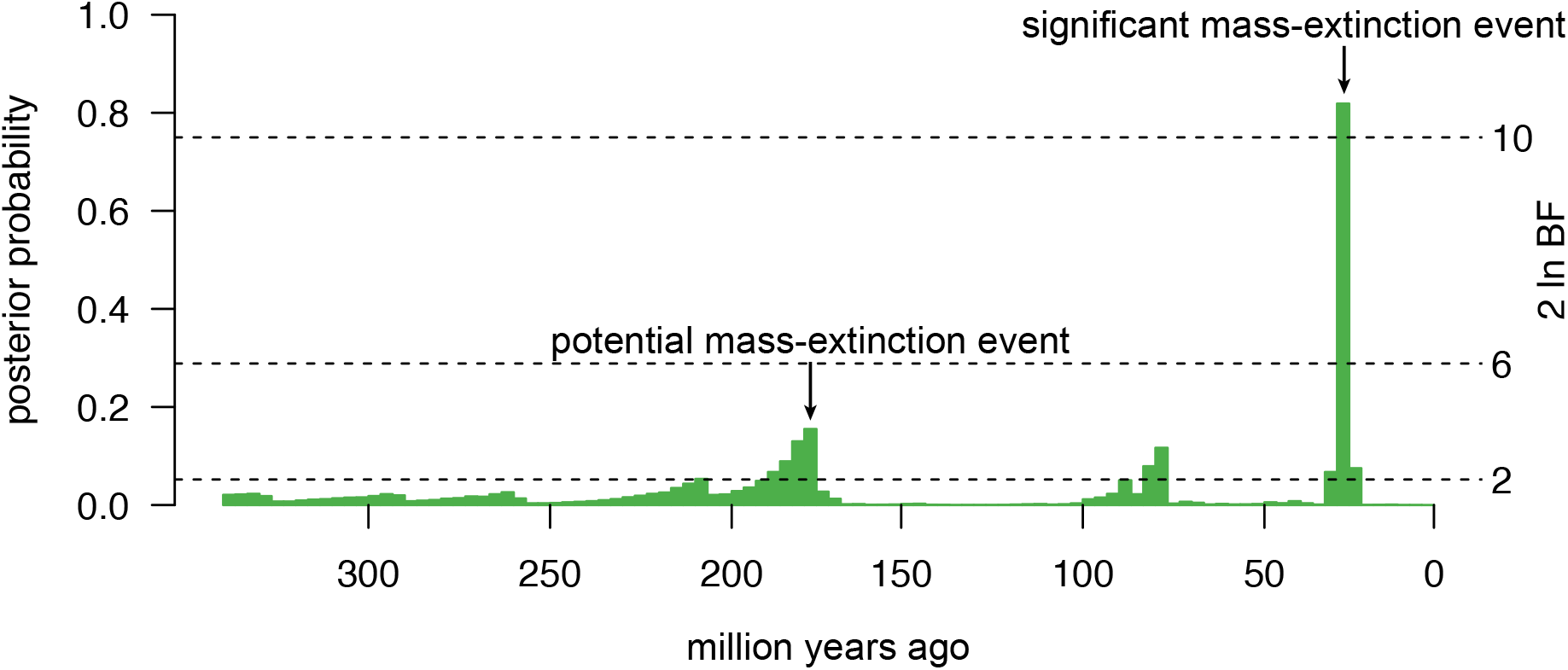
Mass-extinction events in conifers. We inferred two mass-extinction events: a recent event at 23 Ma (with very strong support), and an earlier event at 173 Ma (with positive support).

## DISCUSSION

We present a novel Bayesian approach for detecting mass-extinction events from trees inferred from molecular sequence data—the CPP on mass-extinction time (

~~~
CoMET
~~~

) model—and describe the behavior of our method via a comprehensive simulation study. Simulation studies are critical both for validating statistical methods—to ensure that the algorithms and implementation provide reliable estimates under controlled conditions—and also for providing practical advice for the application of these methods to empirical data. Overall, our simulation study reveals that the 

~~~
CoMET
~~~

 model is statistically well behaved: the method has substantial power to detect the number of mass-extinction events, provides precise and unbiased estimates of the timing of mass-extinction events, and exhibits an appropriate false discovery rate even when background diversification rates may vary. Below, we first consider the implications of our simulation study for the practical application of the 

~~~
CoMET
~~~

 model, and then discuss various avenues for usefully extending the model.

### Practical Application of the CoMET Model

#### Specifying (hyper)priors of the CoMET model

The perennial issue of prior specification is particularly acute for the 

~~~
CoMET
~~~

 model for two reasons. First, our approach relies on the CPP model to infer the history of episodic events—tree-wide shifts in diversification rate and mass-extinction events—and this model is known to be sensitive to the choice of priors. Second, there is typically little biological basis for specifying priors for some of the 

~~~
CoMET
~~~

 model parameters, particularly those describing the frequency and magnitude of shifts in speciation and extinction rates. We should therefore be concerned that the choice of poorly specified (hyper)priors may cause inflated false discovery rates (where the method identifies spurious mass-extinction events) and/or biased parameter estimates (where the method systematically over- or under-estimates the timing of inferred mass-extinction events). Fortunately, the results of our simulation study provide clear guidance on this issue.

Our simulation study demonstrates that the 

~~~
CoMET
~~~

 model is relatively robust to (mis)specification of the hyperpriors describing the frequency of events—*i.e.*, the specified values for the Poisson-rate parameters *λ*_𝔹_, *λ*_𝔹_, and *λ*_𝕄_ (*c.f.*, Figures 5, S2, S3). Nevertheless, we recommend performing analyses under a range of event-rate priors to assess their potential impact (as demonstrated in our conifer analyses; *c.f.*, Figure S9). Although the 

~~~
CoMET
~~~

 model appears quite robust to the specified hyperpriors on event *frequencies*, our simulation reveals that it is somewhat sensitive to the hyperpriors on event *magnitudes*. This is unproblematic for the focal parameter, as we will typically have ample evidence regarding the expected magnitude of mass-extinction events (*i.e.*, to specify values for the *α* and *β* hyperpriors describing the shape of the beta prior on survival probability). By contrast, we will typically lack information regarding the expected magnitude of diversification-rate shifts—*i.e.*, to specify values of the hyperpriors describing the shape of the lognormal speciation- and extinction-rate priors (*μ*_𝔹_, *σ*_𝔹_, and *μ*_𝔹_, *σ*_𝔹_, respectively).

Under simulation, very poorly specified diversification-rate hyperpriors caused the overall false discovery rate of the 

~~~
CoMET
~~~

 model to become slightly elevated: we detected spurious mass-extinction events in 8.1% or 9.9% of the trees simulated under constant or episodically shifting background diversification rates, respectively (Figures S2 and S3). Fortunately, our simulation study indicates that the empirical Bayesian procedure provides a reliable means for specifying appropriate values for the diversification-rate hyperpriors. Under this approach, the false discovery rate of the 

~~~
CoMET
~~~

 model was slightly conservative (we detected spurious mass-extinction events in 2.8% or 4.1% of the trees simulated under constant or episodically shifting background diversification rates, respectively). Accordingly, we strongly recommend use of the empirical Bayesian approach for specifying diversification-rate hyperpriors of the 

~~~
CoMET
~~~

 model (in fact, we have automated this procedure and made it the default setting in our 

~~~
TESS
~~~

 software package).

#### Size and relative age of empirical trees

Our simulation provides guidance on key aspects of study trees that render them appropriate candidates for analysis using the 

~~~
CoMET
~~~

 model. The power of the 

~~~
CoMET
~~~

 model to detect mass-extinction events increases with tree size: all else being equal, it will therefore be easier to detect mass-extinction events in larger trees. Nevertheless, detection rates were quite high even for the smallest trees in our simulation (with *N* = 100 species). Although size is clearly an important factor, the *age* of the tree relative to the mass-extinction time has a much greater impact on the detection probability. In order for a mass-extinction event to leave a detectable phylogenetic signal, the study tree must both contain a su?cient number of lineages at the time of the event, and must also be afforded a sufficient recovery period following a mass-extinction event. Therefore, a mass-extinction event that occurs too close to the root of the study tree will be difficult to detect because too few lineages will have participated in that event (see Figure 6). Similarly, a mass-extinction event that occurs too close to the tips of a study tree will be difficult to detect because too little time has elapsed for the tree to recover from the event. Accordingly, there is a ‘sweet spot’ where our ability to detect a mass-extinction event will be greatest: a candidate study tree should ideally be *∼* 2 − 3 times the age of the putative mass-extinction event, but the event should not occur in the last *∼* 15% of the tree height. This same reasoning explains the power of the 

~~~
CoMET
~~~

 model to detect sequential mass-extinction events. An earlier mass-extinction event has to occur sufficiently far from the root of the tree to ensure that an adequate number of lineages are exposed to the event, but it must also have sufficient time to recover from that event before a subsequent mass-extinction event occurs.

### Future Extensions of the CoMET Model

The 

~~~
CoMET
~~~

 model describes the history of three types of events: tree-wide shifts in speciation rate, tree-wide shifts in extinction rate, and mass-extinction events. Our focus here, however, is restricted to inferring the number and timing of mass-extinction events. Accordingly, we have explicitly adopted the perspective that episodic changes in diversification rate are merely ‘nuisance’ parameters of the 

~~~
CoMET
~~~

 model. These events are primarily included because diversification-rate shifts are thought to be a common feature of empirical trees that, if ignored, might impact our estimates of the focal model parameters. Of course, it might also be possible to use the 

~~~
CoMET
~~~

 model to infer the number, timing, and magnitude of tree-wide shifts in diversification rate. However, we have not investigated the ability of the 

~~~
CoMET
~~~

 model to provide reliable estimates of tree-wide diversification-rate shifts, and so caution users against over-interpreting estimates of these parameters.

The 

~~~
CoMET
~~~

 model is currently restricted to the analysis of trees that have a complete—or if incomplete, a random sample—of species. Given that most empirical trees include a non-random subsample of species, this may limit the application of our method. Of course, it is possible to apply the 

~~~
CoMET
~~~

 model to trees with non-random species sampling, although the statistical behavior of the method is unknown for these datasets. Instead, it would be preferable to provide greater flexibility for accommodating incomplete and non-uniform taxon sampling. For example, we could explicitly incorporate various departures from random species sampling—such as diversified species sampling (*c.f.*, Höhna et al. 2011; Cusimano et al. 2012; Höhna 2014)—in the 

~~~
CoMET
~~~

 model. In fact, the presented likelihood equations of the episodic birth-death process with explicit mass-extinction events and the Bayesian inference framework can readily be applied to diversified species sampling (Höhna 2014; 2015). This is an area of current work (Höhna, May, and Moore, *in prep.*).

Currently, the 

~~~
CoMET
~~~

 model assumes that a mass-extinction event is equally likely to occur at any time over the interval spanned by the study tree. Our use of a uniform prior on mass-extinction times is therefore somewhat naive, as this expectation effectively ignores relevant information regarding the probable timing of mass-extinction events. Fortunately, a straightforward extension of the 

~~~
CoMET
~~~

 model would allow the use of informative priors that reflect our knowledge regarding probable mass-extinction times. Similarly, the 

~~~
CoMET
~~~

 model presently assumes that a mass-extinction event is equally likely to impact all contemporaneous lineages in the study tree. Our use of a uniform prior on the survival probability across lineages may also be somewhat naive. Imagine, for example, that other variables might render species more or less susceptible to mass-extinction events. This possibility could be addressed by extending the 

~~~
CoMET
~~~

 model to allow the survival probability of a lineage to depend on the inferred state of a continuous (*e.g.*, body size, metabolic rate, range size) or a discrete (*e.g.*, marine/terrestrial, endothermic/ectothermic) variable.

As currently implemented, the 

~~~
CoMET
~~~

 model effectively treats the study tree as an observation. Phylogenies are, of course, inferences from data, and so entail (sometimes considerable) uncertainty. Ignoring this phylogenetic uncertainty will therefore tend to make us overly confident in our conclusions regarding mass-extinction events. We could accommodate phylogenetic uncertainty by extending the 

~~~
CoMET
~~~

 model in one of two ways. A *sequential Bayesian* approach provides a simple (albeit computationally intensive) solution: mass-extinction events could simply be inferred by integrating over a posterior distribution of trees that has previously been estimated using some other program. Alternatively, a *hierarchical Bayesian* approach involves jointly inferring the phylogeny, divergence times, and history of mass-extinction events. This solution would require considerably more effort, as it would require implementation of the 

~~~
CoMET
~~~

 model within an existing Bayesian phylogenetic inference program, such as RevBayes (Höhna et al. 2015). Although more involved, this would provide an elegant solution for accommodating phylogenetic uncertainty that would permit more robust inferences regarding mass-extinction events.

Currently, the 

~~~
CoMET
~~~

 model is limited to the analysis of a single study phylogeny. However, we may also want to explore the impact of mass-extinction events on a set of trees. We might, for example, wish to study the effect of mass extinction on the flora of a particular geographic region that is comprised of several distantly related plant groups. This inference scenario could be addressed by extending our approach by allowing the parameters of the 

~~~
CoMET
~~~

 model to be inferred from a composite vector of waiting times for a set of trees. Moreover, we could either assume that the survival probability is identical for all groups, or allow each group to have a unique response to episodes of mass extinction. This extension is relatively straightforward, and would simultaneously extend the range of questions that can be addressed using the 

~~~
CoMET
~~~

 model, and also increase the power of the method to detect mass-extinction events by virtue of increasing the effective tree size.

## SUMMARY

We present a novel Bayesian approach—the 

~~~
CoMET
~~~

 model—that provides an effective tool for identifying mass-extinction events in molecular phylogenies, even when those groups have experienced more prosaic temporal variation in diversification rates. We performed a thorough simulation study to characterize the statistical behavior of this new approach, which reveals that the 

~~~
CoMET
~~~

 model has substantial power to detect the number of mass-extinction events, provides precise and unbiased estimates of the timing of mass-extinction events, and exhibits an appropriate false discovery rate. Based on the results of our simulation study, we offer some practical advice for applying the method to empirical datasets—including guidance regarding the choice of (hyper)priors, and insights on the properties of study trees that will impact detection probabilities using our method. We also demonstrate the empirical application of the 

~~~
CoMET
~~~

 model to a recent phylogeny of conifers, which reveals that this group experienced two major episodes of mass extinction. We are optimistic that the development of a robust and powerful statistical approach for detecting mass-extinction events will provide an important tool for advancing our understanding of how events in Earth history have shaped the Tree of Life.

## Acknowledgements

We are grateful to John Huelsenbeck, Bruce Rannala, and Michael Donoghue for thoughtful discussion, and to Michael Donoghue for providing access to the conifer dataset. This research was supported by National Science Foundation (NSF) grants DEB-0842181, DEB-0919529, DBI-1356737, and DEB-1457835 awarded to BRM. Computational resources for this work were provided by NSF XSEDE grants DEB-120031, TG-DEB140025, and TG-BIO140014 awarded to BRM. SH was funded by the Miller Institute for Basic Research in Science.

## APPENDIX

Here we provide a single equation to calculate the probability of an observed tree. For convenience of notation, we construct a unique vector, 𝕏, that contains all of the divergence times and event times (for shifts in speciation and extinction rates, and mass-extinction events) sorted in **increasing** order (see Fig. 1.b). It is convenient to expand the vectors for all of the other parameters so that they have the same number of elements as 𝕏. We use the notation *S*(2*, t*_1_ = 0*, T*) to represent the survival of two lineages in the interval [*t*_1_*, T*], which is the condition we enforce on the reconstructed evolutionary process. This allows us to write the more convenient equation for the probability density of a reconstructed tree

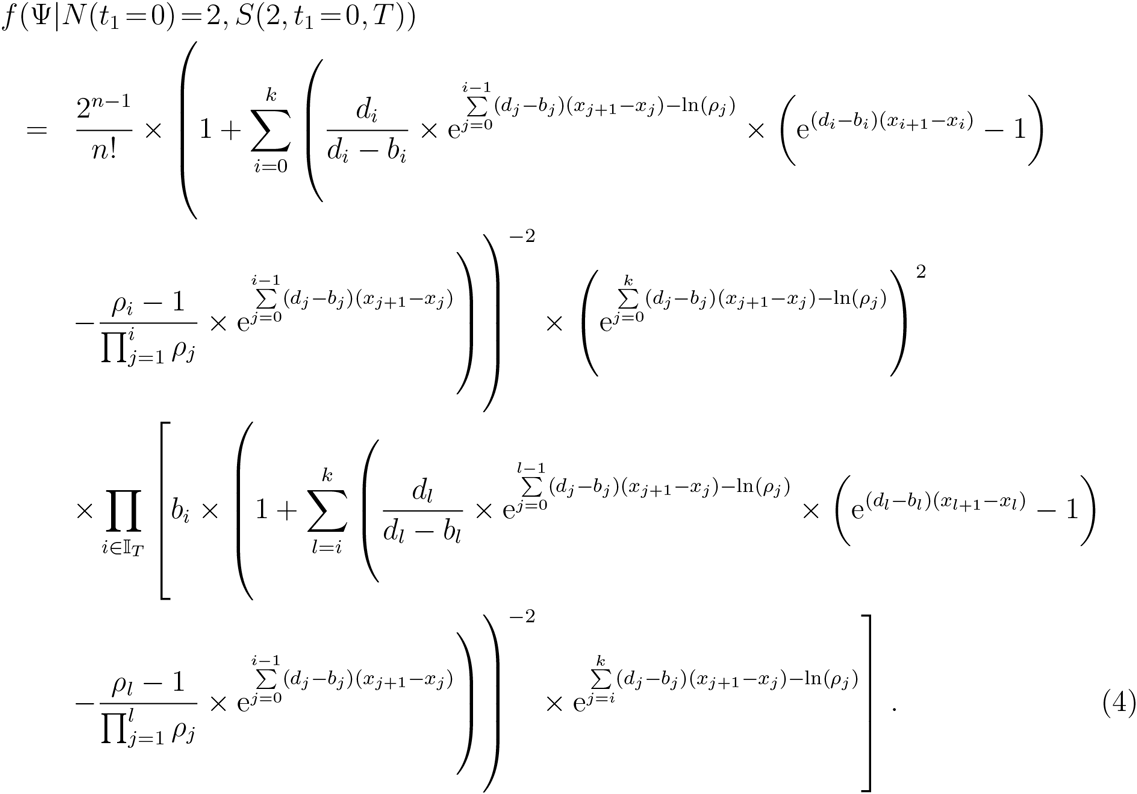

Additional details regarding the derivation of this probability density and its relation to other birth-death models are discussed by Höhna (2015).

## Supporting Information

### CoMET: A Graphical Model Description

Below, we provide a graphical-model representation of the 

~~~
CoMET
~~~

 model (Figure S1). The phylogenetic application of graphical models is described by Höhna et al. (2014).

**Figure S1:**
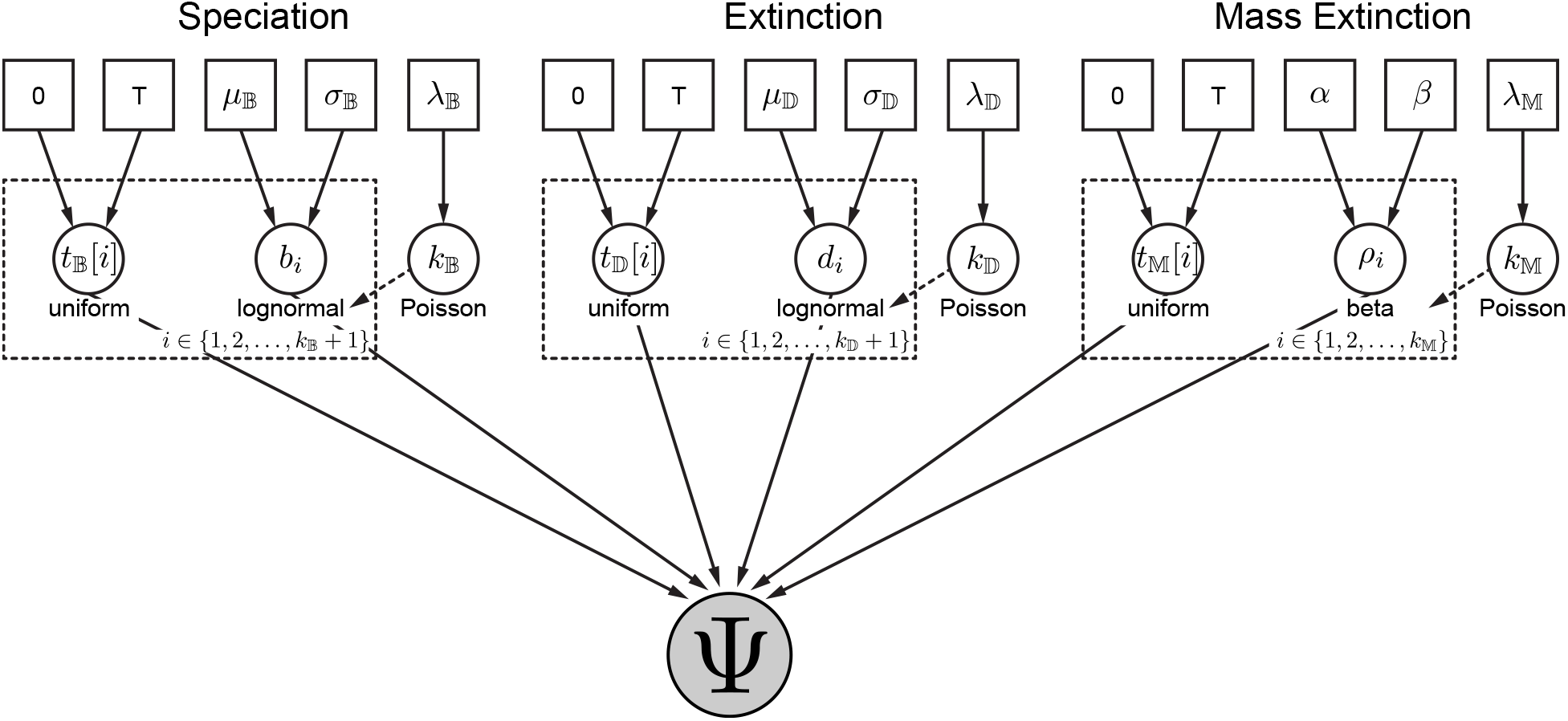
The CoMET model graph. By convention, *constant parameters*—such as the rate of the Poisson process, *λ*_*i*_—are enclosed in solid squares, whereas *random variables*— such as the number of speciation-rate shifts, *k*_𝔹_, or the number of mass-extinction events, *k*_𝕄_—are enclosed in solid circles, and the observations—the vector of waiting times in the tree, ψ—are enclosed in a shaded circle to indicate that these variables have been observed. Here the choice of prior distribution and the hierarchical structure of the model is explicit; the number of speciation-rate shifts, extinction-rate shifts, and mass-extinction events are Poisson distributed variables—*k*_𝔹_, *k*_𝔹_, and *k*_𝕄_, respectively—where lognormal distributions are used for the speciation and extinction rates and a beta distribution is used for the survival probability. Arrows indicate the dependence between parameters, where the direction specifies the conditional relation (*e.g.,* an arrow from the constant parameters 0 and *T* to the random variable *t*_𝔹_[*i*] indicates that *t*_𝔹_[*i*] is conditional on 0 and *T*). Dashed squares (‘plates’) indicate repetition; here the number of replicates depends on the variables *k*_𝔹_, *k*_𝔹_, and *k*_𝕄_.

**Figure S2:**
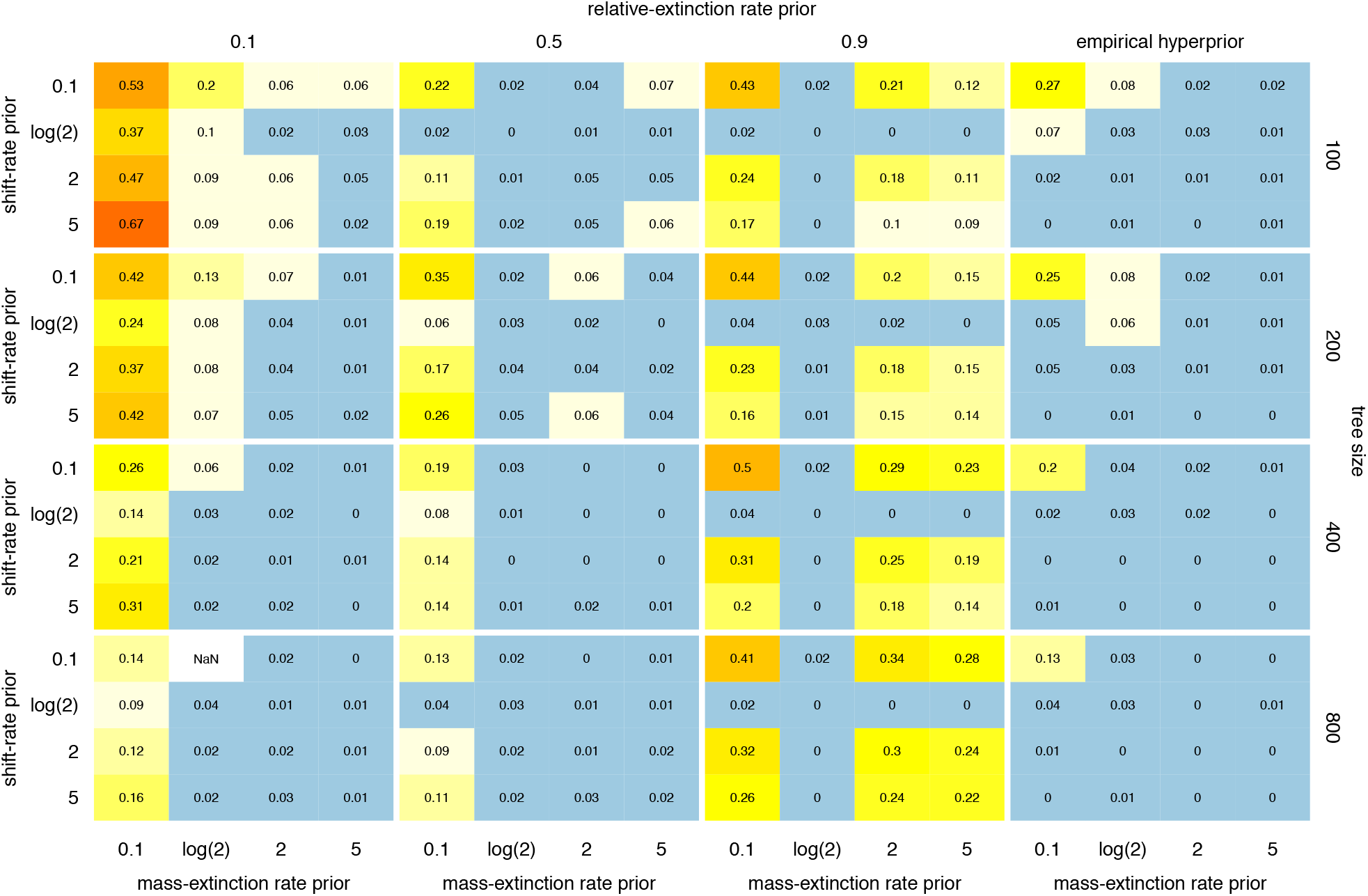
False discovery rates. Frequency of detecting spurious mass-extinction events in the absence of background shifts in diversification rates. Rows of panels correspond to different tree sizes (*N* = {100, 200, 400, 800}), and columns of panels correspond to different priors on the relative extinction rate (*μ*_𝔻_ = 0.1, 0.5, 0.9, empirical). Within each panel, the rows correspond to false discovery rates under various priors on the number of diversification rate shifts (rows) and mass extinction events (columns).

**Figure S3:**
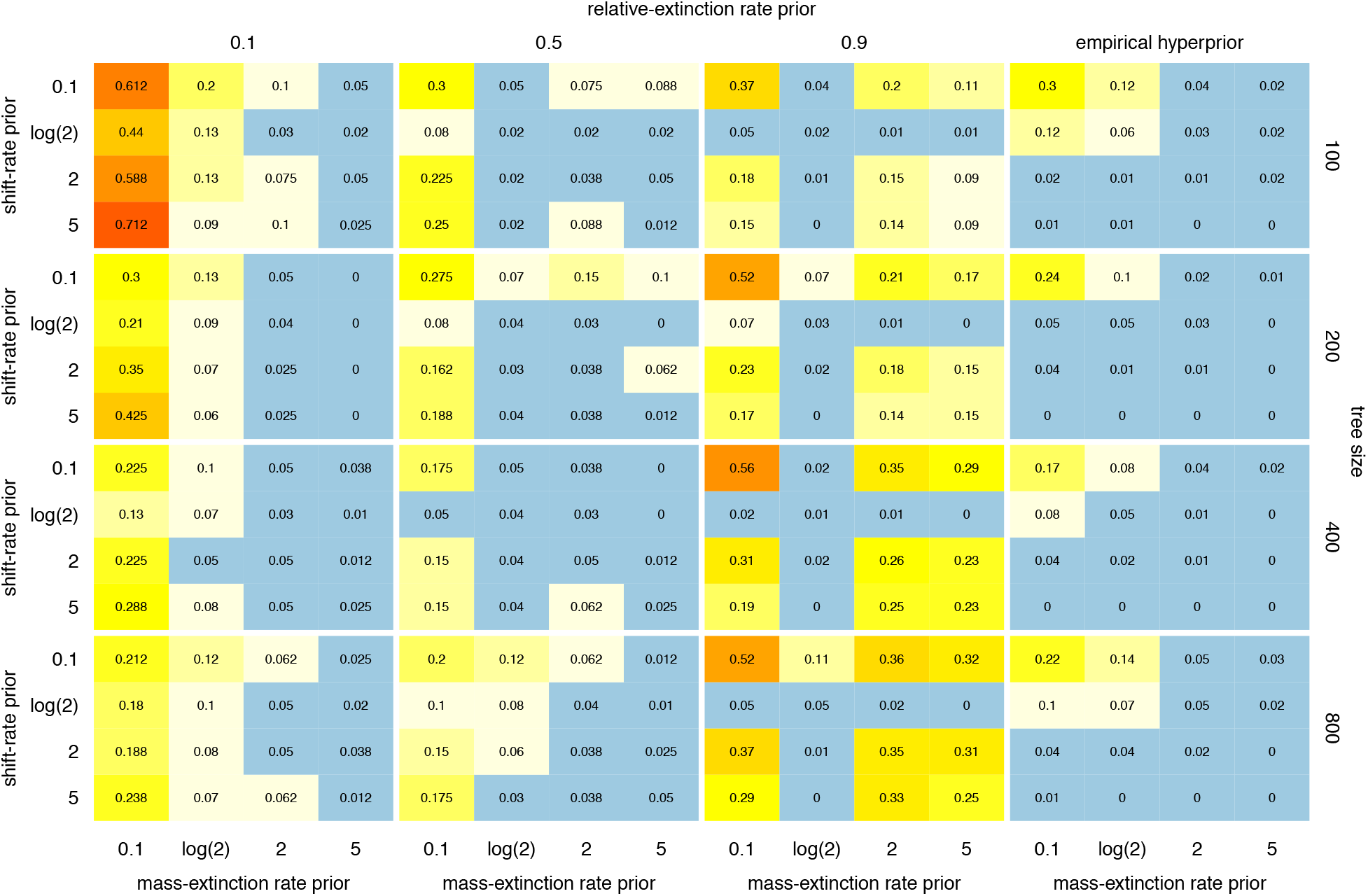
False discovery rates. Frequency of detecting spurious mass-extinction events in the presence of background shifts in diversification rates. Rows of panels correspond to different tree sizes (*N* = {100, 200, 400, 800}), and columns of panels correspond to different priors on the relative extinction rate (*μ*_𝔻_ = 0.1, 0.5, 0.9, empirical). Within each panel, the rows correspond to false discovery rates under various priors on the number of diversification rate shifts (rows) and mass extinction events (columns).

**Figure S4:**
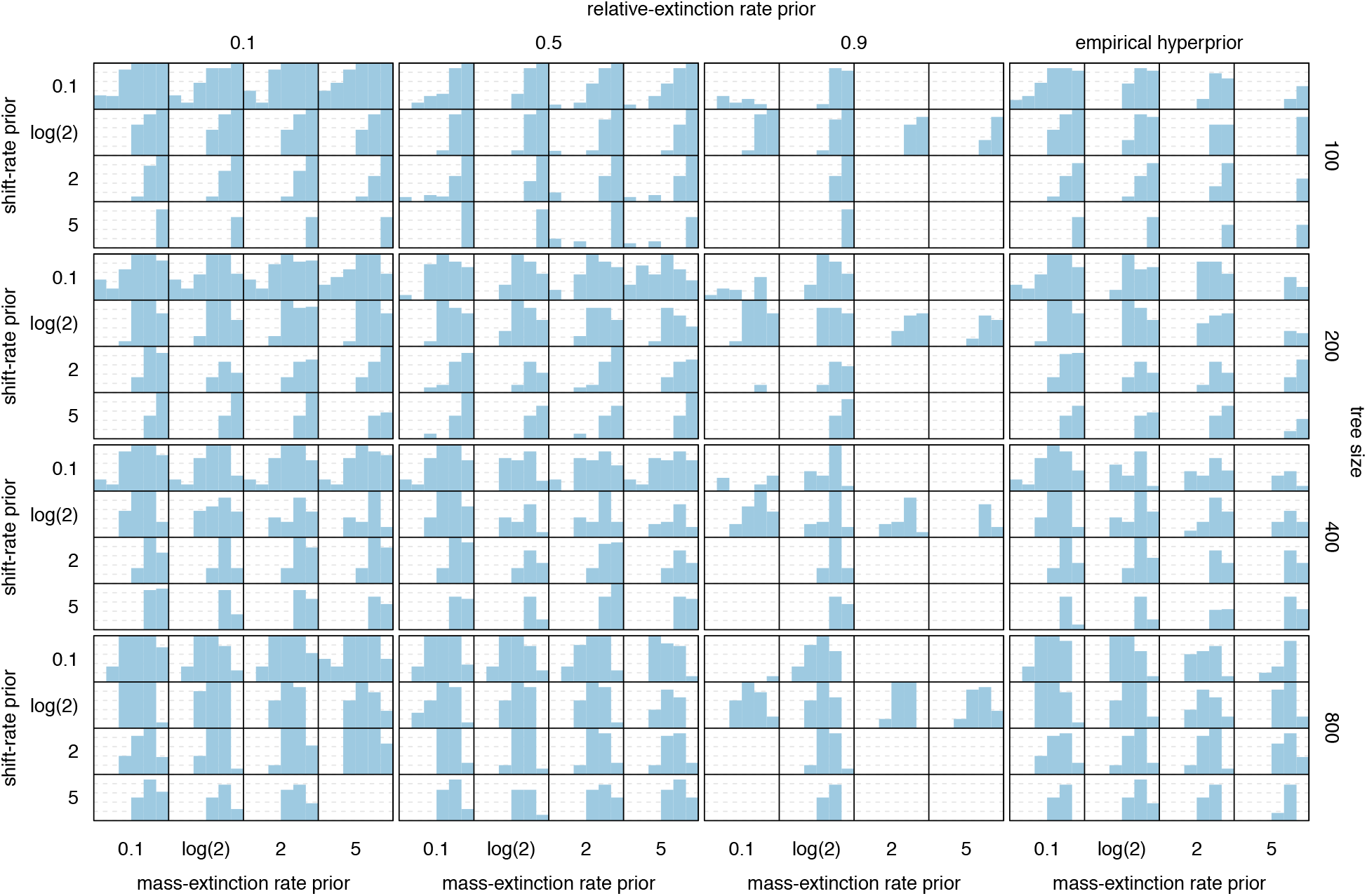
Power as a function of time. Frequency of correctly identifying mass-extinction events in the absence of background shifts in diversification rates. Rows of panels correspond to different tree sizes (*N* = {100, 200, 400, 800}), and columns of panels correspond to different priors on the relative extinction rate (*μ*_𝔻_ = 0.1, 0.5, 0.9, empirical). Within each panel, the rows correspond to the power under various priors on the number of diversification rate shifts (rows) and mass extinction events (columns). In each cell, we compute the power as a function of time by binning simulated trees into the interval corresponding to their mass-extinction time, and computing the fraction of those trees where a mass-extinction event was correctly inferred.

**Figure S5:**
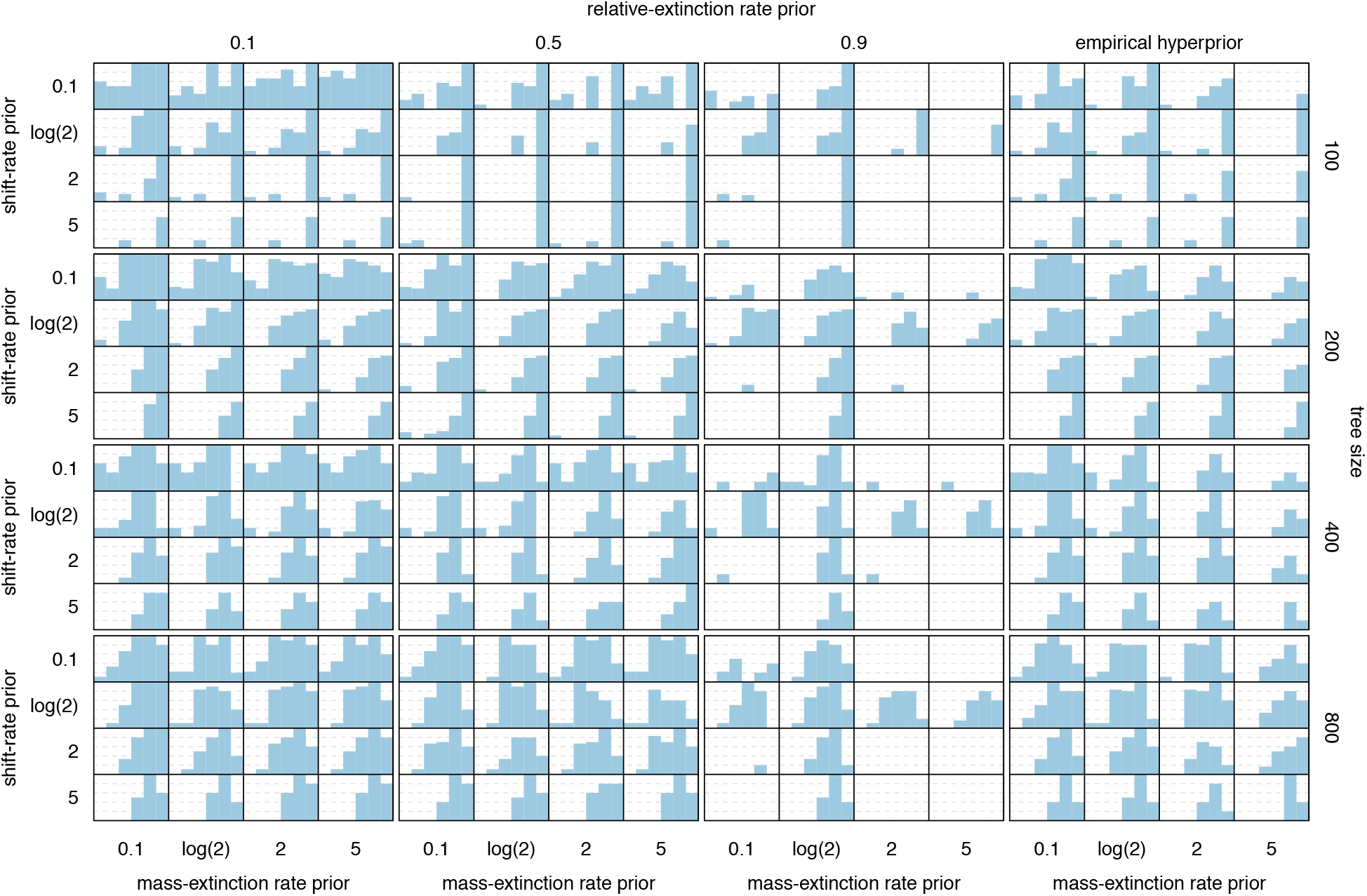
Power as a function of time. Frequency of correctly identifying mass-extinction events in the presence of background shifts in diversification rates. Rows of panels correspond to different tree sizes (*N* = {100, 200, 400, 800}), and columns of panels correspond to different priors on the relative extinction rate (*μ*_𝔻_ = 0.1, 0.5, 0.9, empirical). Within each panel, the rows correspond to the power under various priors on the number of diversification rate shifts (rows) and mass extinction events (columns). In each cell, we compute the power as a function of time by binning simulated trees into the interval corresponding to their mass-extinction time, and computing the fraction of those trees where a mass-extinction event was correctly inferred.

**Figure S6:**
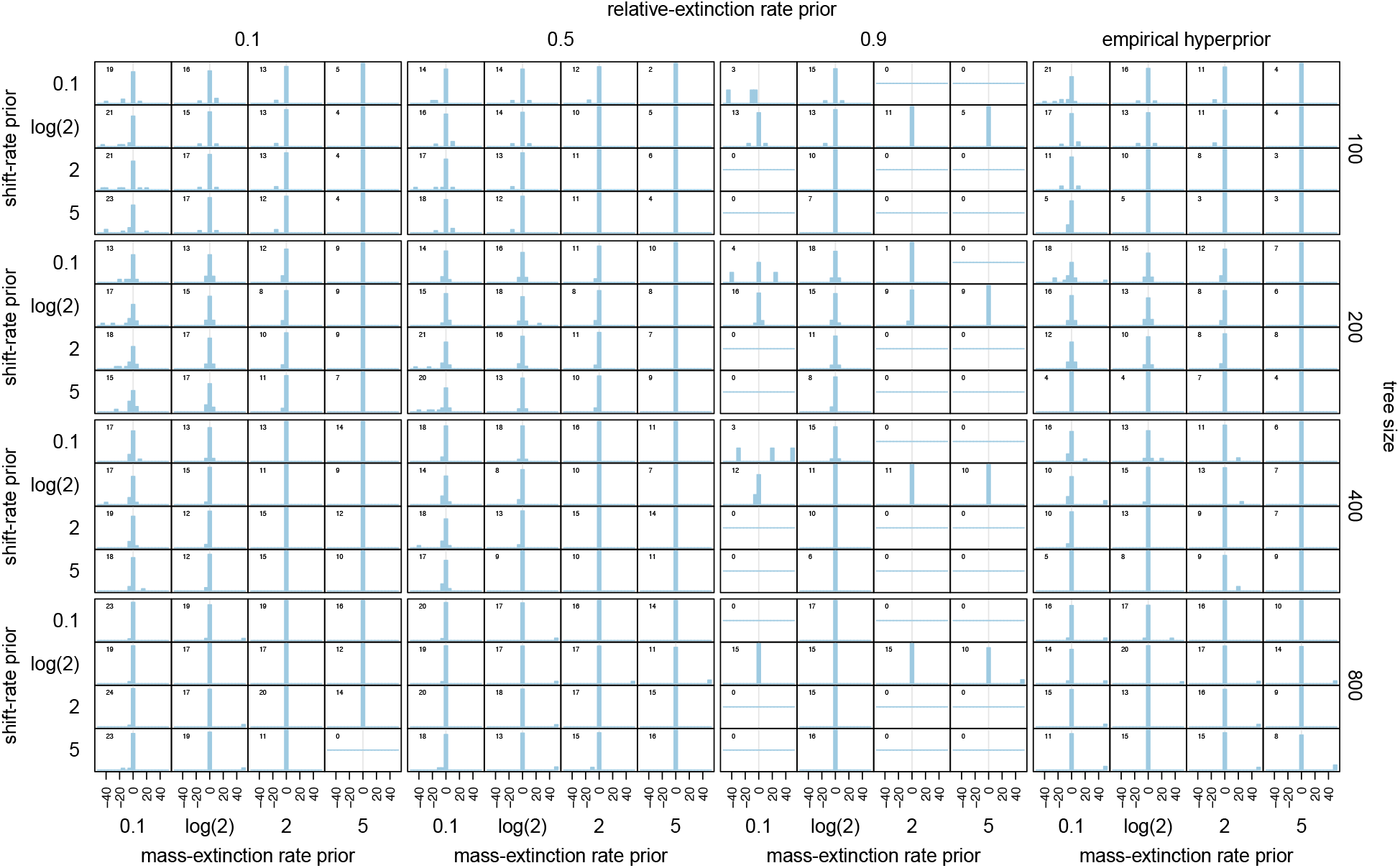
Bias in estimates of mass-extinction time. The distribution of bias in estimated mass-extinction times in the absence of background shifts in diversification rates. Rows of panels correspond to different tree sizes (*N* = {100, 200, 400, 800}), and columns of panels correspond to different priors on the relative extinction rate (*μ*_𝔻_ = 0.1, 0.5, 0.9, empirical). Within each panel, the rows correspond to the bias under various priors on the number of diversification rate shifts (rows) and mass extinction events (columns). The bias is computed as (*t*_simulated event_ − *t*_estimated event_)*/*tree height *×* 100%.

**Figure S7:**
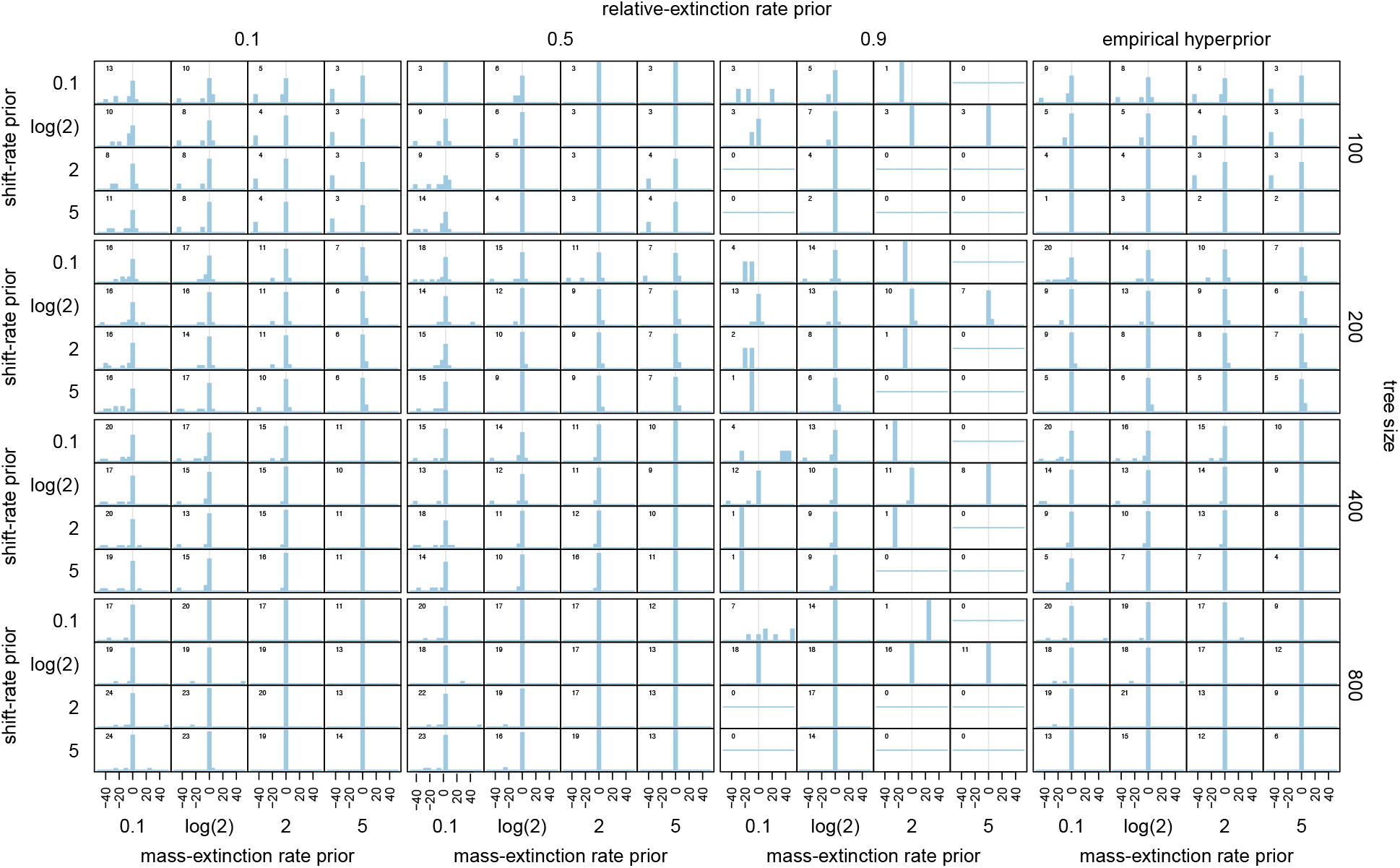
Bias in estimates of mass extinction time. The distribution of bias in estimated mass-extinction times in the presence of background shifts in diversification rates. Rows of panels correspond to different tree sizes (*N* = {100, 200, 400, 800}), and columns of panels correspond to different priors on the relative extinction rate (*μ*_𝔻_ = 0.1, 0.5, 0.9, empirical). Within each panel, the rows correspond to the bias under various priors on the number of diversification rate shifts (rows) and mass extinction events (columns). The bias is computed as (*t*_simulated event_ − *t*_estimated event_)*/*tree height *×* 100%.

**Figure S8:**
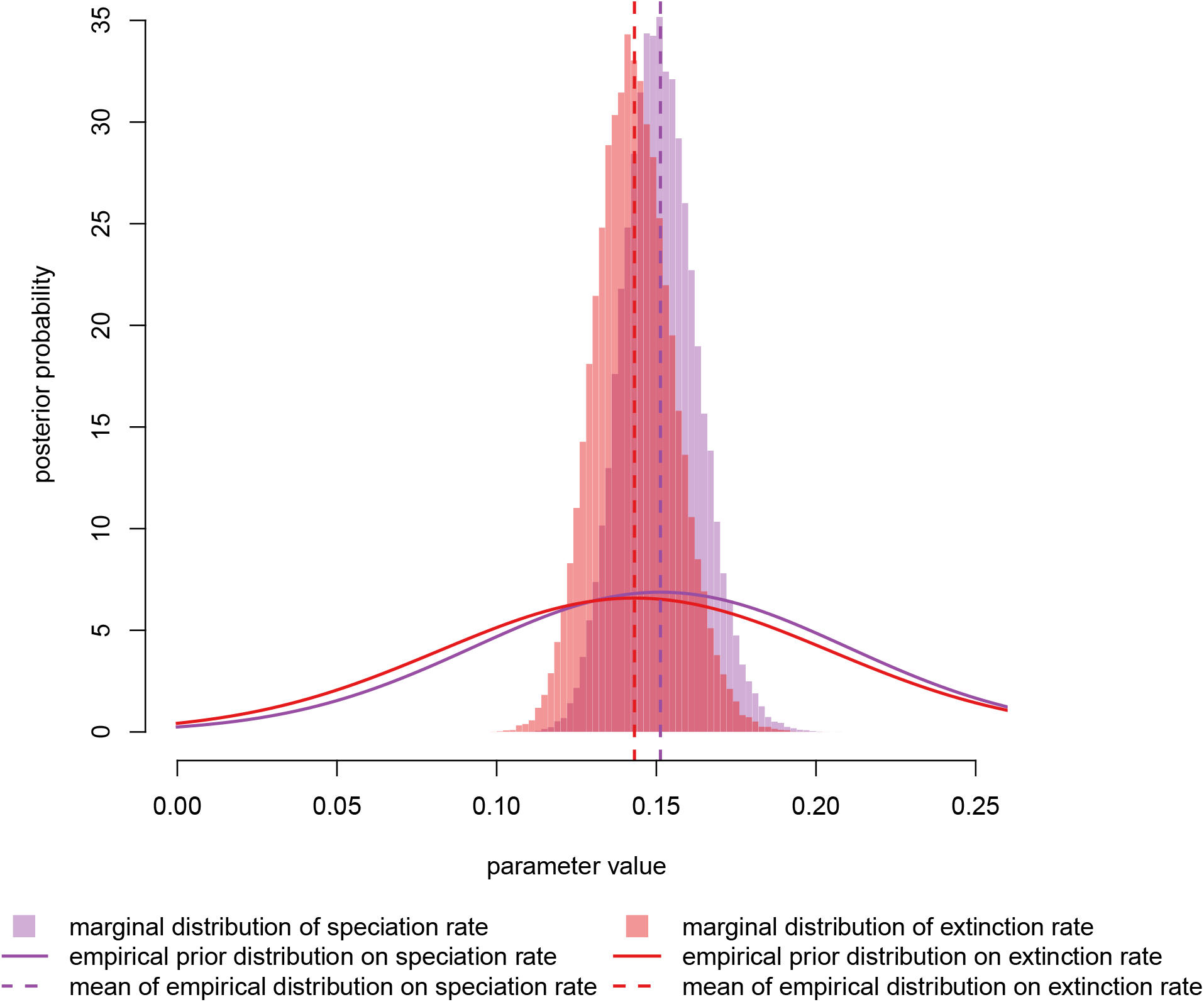
Empirical hyperpior analysis of the conifers. Histograms are the marginal posterior densities of the speciation (purple) and extinction (red) rates for the constant-rate birth-death-sampling process applied to the conifer data. Solid lines are the corresponding marginal prior densities of the speciation and extinction rates used for the subsequent 

~~~
CoMET
~~~

 analyses, dashed vertical lines are means of the empirical prior densities.

**Figure S9:**
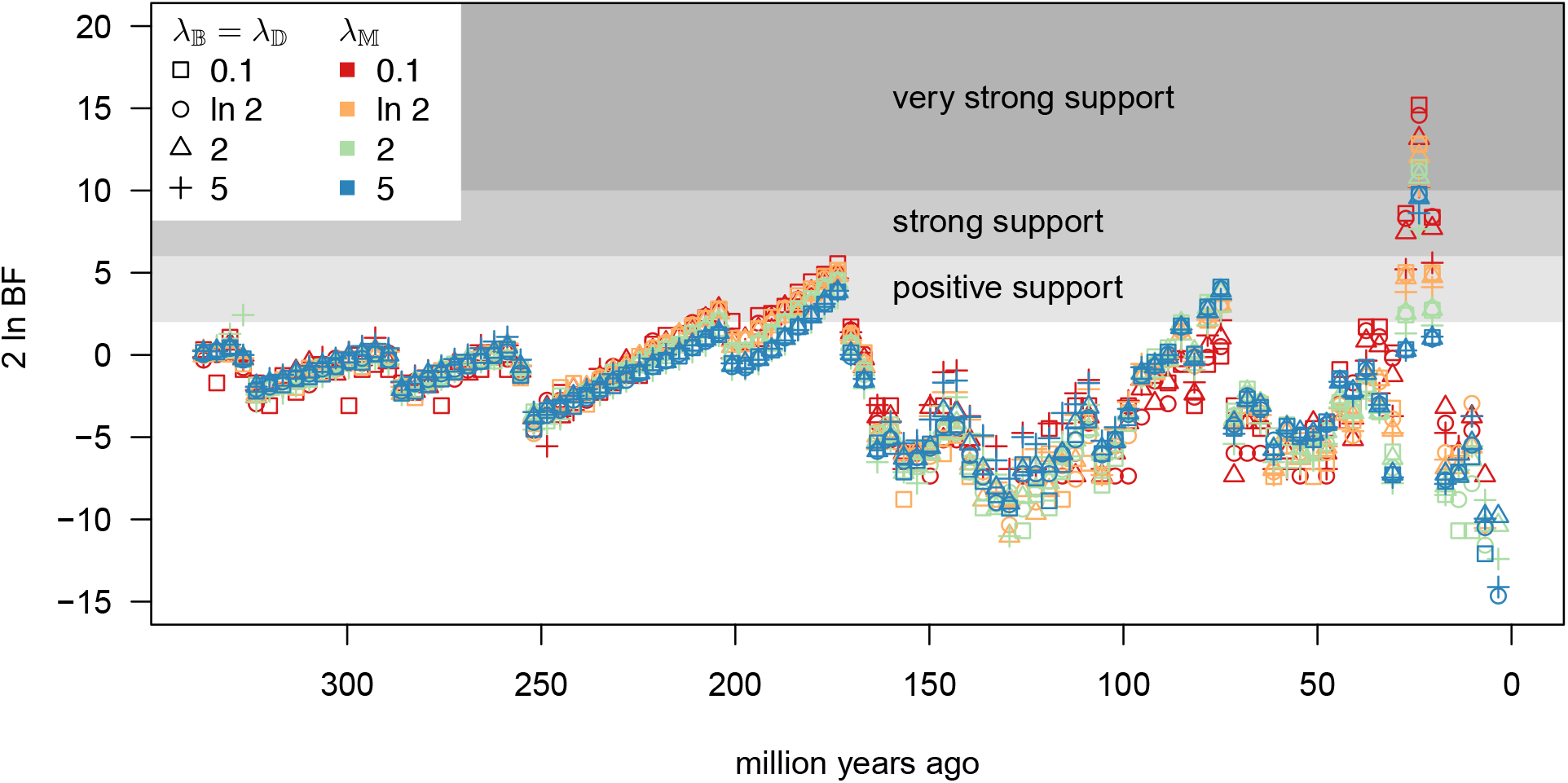
Bayes factors for conifer analyses under various priors. We divided the time period (0, 340.43) into 100 discrete intervals and computed the Bayes factor support for there being at least one mass-extinction event in each interval. Point types and colors correspond to different combinations of prior settings on *λ*_𝔹_*, λ*_𝔹_, and *λ*_𝕄_ (see legend). Bayes factor support is fairly consistent across all the prior settings; in particular, there is always at least strong support for a mass-extinction event about 23 Ma, and consitently high positive support for another mass-extinction event at approximately 173 Ma.

